# Expanding single-cell toolbox with a cost-effective full-length total RNA droplet-based sequencing technology

**DOI:** 10.1101/2025.02.23.639726

**Authors:** Xue Sun, Shir-Liya Dadon, Dena Ennis, Wenpeng Fan, Muhammad Awawdy, Eli Reuveni, Adi Alajem, Oren Ram

## Abstract

sc-rDSeq is a scalable, full-length total RNA droplet-based technology that captures both polyadenylated and non-polyadenylated RNAs, including histone-RNAs, small and long non-coding RNAs, and enhancer RNAs. It achieves a tenfold increase in UMIs per cell over inDrops while remaining simple and cost-efficient. Applied to lung cancer cells, sc-rDSeq revealed hidden heterogeneity, divergent signaling pathways, and non-polyA RNA variations undetectable with 3’ end-based methods. In EGFR inhibitor-treated persister cells, it confirmed drug-induced cell cycle arrest based on histone mRNA expression, identified distinct resistance strategies. It also detected alternative splicing events and enabled single cell genotyping of SNVs linked to drug resistance. While SNVs act as drivers of drug resistance, additional layers of expression-based cellular states must be explored to fully understand and combat persister cells. These findings highlight the potential of sc-rDSeq to advance transcriptomic studies and personalized medicine by providing a comprehensive and scalable approach to cellular RNA analysis.

## INTRODUCTION

Over the last decade, single-cell RNA-sequencing (scRNA-seq) has become an essential tool for understanding cellular heterogeneity in complex biological systems such as tissue development and tumor environments^1,2^. Significant efforts have been made to advance the technical aspects of scRNA-seq technology. Just eight years ago, only a few hundred cells could be sequenced at a time, but today, hundreds of thousands of single cells can be sequenced with greater sensitivity, enabling accurate characterization of cell types, states, and unique populations. However, most methods rely on oligo-dT priming, which captures only polyadenylated RNA (polyA+) and are limited to the length of reads with Unique Molecular Identifiers (UMIs) for PCR deduplication^3^, which covers up to 600bp upstream to the 3’ end, or downstream to the 5’ end^4,5^. As a result, non-polyadenylated RNA (polyA-), including non-coding RNA (ncRNA), histone RNA, micro-RNA, and enhancer RNA (eRNA), and alternative RNA isoforms remains undetected. This limitation restricts our ability to study these crucial transcriptome fractions and prevents us from exploring key transcriptional features such as alternative splicing events and, potentially, the detection of RNA-based mutations.

Currently, the most widely used single-cell full-length profiling methods are Smart-Seq2 and Smart-Seq3^6,7^. However, they are limited to polyA+ RNAs and are cost-prohibitive due to the use of Tn5 transposase and well-plate-based single-cell isolation steps. Several other single cell full-length RNA sequencing methods have been developed and shown to have a decent coverage over total RNA. RamDA-seq^8^ utilizes Not-So-Random (NSR) and polyT primers together for capturing non-ribosomal RNA. The single cell isolation is based on Fluidigm C1 microfluidics chips, which has limited throughput, and the method does not incorporate UMIs for PCR deduplication. More recent methods Smart-seq-total^9^ and VASA-seq^10^ are based on the polyadenylation of total RNAs, allowing the use of conventional polyT primers for capturing. VASA-seq is currently the only scRNA-seq method capable of capturing full-length total RNA while incorporating UMIs. However, this technology requires two microfluidic steps, making it inaccessible for most laboratories and incompatible with widely available commercial platforms like 10X Genomics. In the first step, cells are captured, and RNA is fragmented with polyA tails added to the fragments. In the second step, reverse transcriptase and primers are introduced, enabling the incorporation of Unique Fragment Identifiers (UFIs) into each RNA fragment. Both capture total RNA, including rRNA, which is later removed during library preparation after amplification, either using rRNA-probes with beads^11^ or with Cas9 targeting rRNA^12^. Since rRNA depletion occurred downstream of the original RNA capture, the rRNA are also amplified, which originally constitute more than 80 - 90% of the total RNA by mass in a cell^13^. This reduces the accessibility of the resources for the amplification of non-rRNA sequences. Consequently, there is a need for a high-throughput, easy-to-use and cost-effective scRNA-seq method for robust detection of full-length total RNA.

Here, we developed and demonstrated sc-rDSeq, a droplet-based ribosomal-depleted sequencing method that captures full-length total RNA. The method is built upon the inDrops^4^, incorporating a major modification in primer design and library preparation. sc-rDSeq is strand-specific, based on linear amplification of *in vitro* transcription, and included UMI for PCR deduplication. rRNA removal is not needed in the library preparation as they are excluded from the initial profiling. Moreover, sc-rDSeq is highly cost-effective for library preparation at the same level as standard inDrops (∼0.08 $/cell). The core of this technology is an easily adaptable set of primers, based on the work of Armour *et al.* ‘s with selective hexamers for cDNA synthesis^14^. Their ‘Not-So-Random’ (NSR) primers significantly reduced rRNA content in bulk comparing to random priming (from 80% to 13%) but were not preformed effectively in droplets (ex. RamDA-seq), due to the significant loss of sensitivity in single-cell profiling and the ability of NSR primers to detect rRNA at low levels^8^. To improve this, we designed a new set of 220 hexamers, ribosomal-depleted sequences (rDS), which efficiently deplete rRNA in drop-based single-cell RNA-seq. Unlike polyT primers, rDS can bind multiple sites on an RNA molecule, enhancing sensitivity and allowing for full-length RNA coverage, including rare RNA species. We first demonstrated the superiority of sc-rDSeq over inDrops in terms of full-length coverage, diverse RNA profiling, and enhanced sensitivity. A Benchmark over cost per cell against other state-of-the-art methods were conducted and we showed the uppermost cost effectiveness of sc-rDSeq.

Non-small cell lung cancer (NSCLC) is one of the most vital cancers in the world. Despite the new generations of drugs, resistance remained occurring in patients^15^. It was believed that genetical mutation is the only driving force for the resistance, however recent studies have demonstrated the diversity of cellular states in response to drug treatment, which suggests more complicated mechanism in transcription regulations^16^. Hence, we decided to leverage sc-rDSeq on studying the heterogeneity of drug-tolerant ‘persister’ (DTP) cells from NSCLC cancer cell line. We profiled more than 7000 DTPs using sc-rDSeq, then characterized the shift in cellular states from naïve to persistent, on both polyA+ and polyA-RNA levels. Notably, G1 arresting is observed in DTPs, supported by both polyA cell cycle marker gene expression and polyA-histone mRNA expression. Canonical pathways were upregulated in DTPs, along with some known maker genes for resistance, while polyA-RNA, like microRNA and other non-coding RNAs were found co-expressed in different cellular clusters. Besides, alternative splicing on pseudo-bulk level and single cell level were also profiled, suggesting an extra dimension of information in understanding the cellular states. Moreover, due to the strand-specificity, regions with enrichment of bi-directionally transcribed RNA were utilized in describing eRNA profiles and showed strong association between eRNA and its targeted gene expression at the single-cell level. Taken together, sc-rDSeq is a cost-effective, high-throughput, and sensitive single-cell method that ensures full-length coverage of total RNA and enhances sensitivity, opening opportunities to explore the interactive information between mRNA and other types of RNA at the single-cell level.

## RESULTS

### sc-rDSeq enables high-throughput detection of total RNA with high sensitivity and full-length coverage

We have developed a high-throughput droplet-based total RNA sequencing method - sc-rDSeq (**Fig. 1a**). In brief, single cells are co-encapsulated with reverse transcription - lysis buffer and hydrogel beads carrying specific barcoded primers allowing for non-ribosomal total RNA capturing. Two types of barcoded primers are employed for achieving this purpose, ribosomal-depleted sequences (rDS) and polyT5C8N primers (**Fig. S1a-c**). rDS primers is a set of 220 hexamers computationally designed and validated to selectively enrich for all non-rRNA targets in cells. In silico alignment revealed that, for every 100 nt of total RNA, 4.4 sequences aligned, capturing 99.9% of total RNA, thus providing ample priming coverage (**Fig. S1d**). We observed that using rDS primers alone results in limited reverse transcription (RT) efficiency. However, when combined with polyT5C8N primers, the number of UMIs captured per cell increases fourfold (**Fig. S2a**). The polyT5C8N primer complements random priming by supporting high efficiency polyT priming while also initiating RT from internal RNA regions. The polyT sequence anneals to polyA repeats, particularly the polyA tail of mRNA, while the CCC sequence acts as a bridge, facilitating the annealing of the random octamer to upstream regions of the mRNA. Notably, since mRNA naturally forms secondary structures, the folded RNA regions are spatially closer to the polyA tail, increasing their accessibility to the 8Ns for annealing (**Fig. S2c**). We integrated the rDS and polyT5C8N primers into single-cell barcode beads, which also included handles for library preparation and Unique Molecular Identifiers (UMIs) to facilitate PCR duplication removal. To improve RT reaction efficiency, we implemented a ramping-cycling-short RT strategy. In this approach, the temperature is gradually ramped from 4°C to 50°C. This ramping program is repeated for 20 cycles to enhance the annealing of initially unbound primers. Additionally, to prevent strand displacement during RT reactions^17^, the total RT reaction time at 50°C is limited to 5 minutes. These modifications significantly improved RT efficiency and increased single-cell complexity from 6k to 116k nUMIs per cell (**Fig. S2b**).

**Fig 1.**
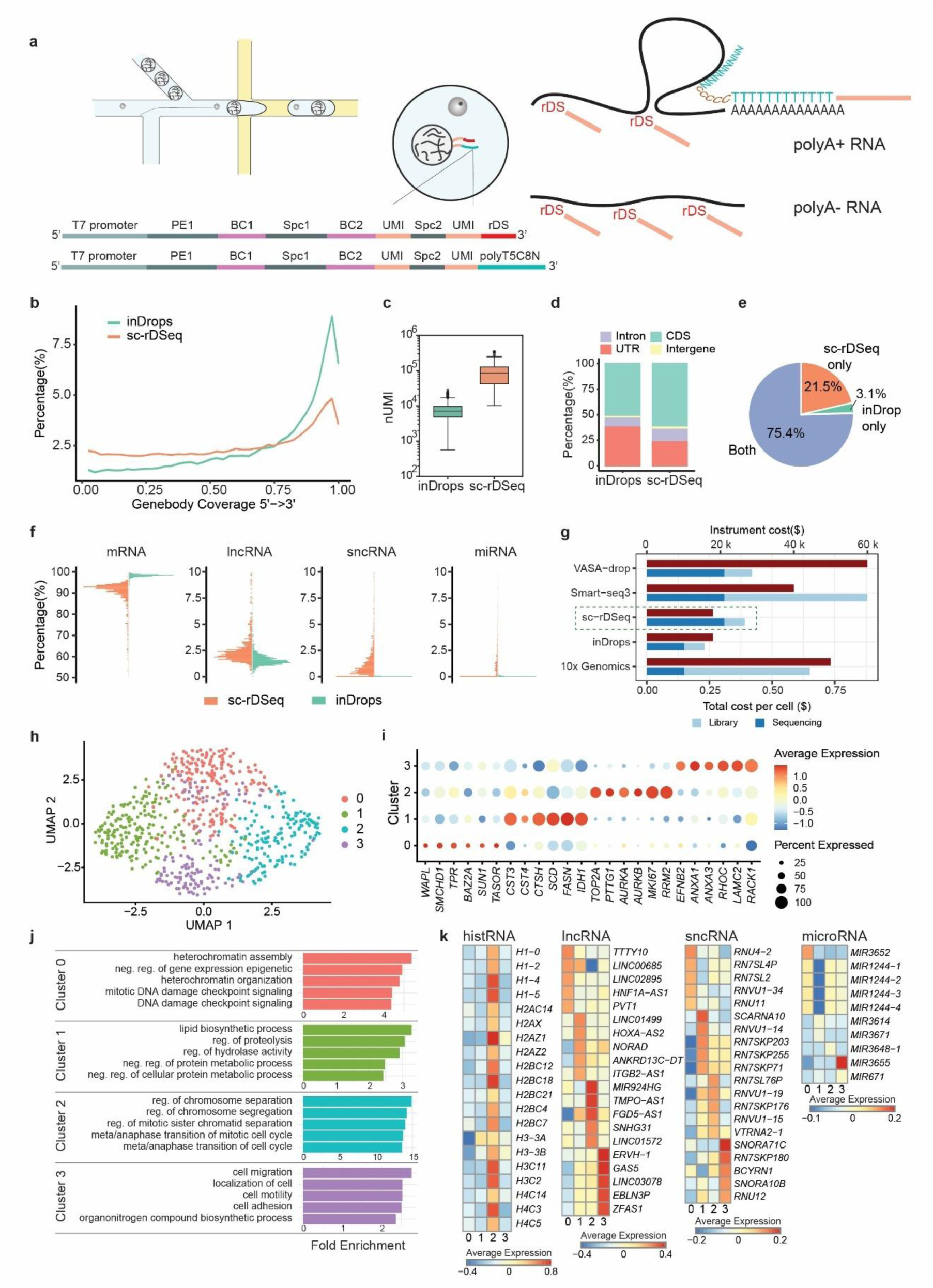
sc-rDSeq showed high sensitivity for full-length total RNA in single cell with high sensitivity. **a)** Single cell and barcode beads are co-encapsulated by microfluidics platform. The primers on the barcode beads are equipped with handles for library preparation and single cell barcode and ended with rDS (red) or polyT5C8N (blue), which target both polyA+ and polyA-RNA transcripts. **b)** Gene-body coverage across all genes based on merged single cells data showing strong 3’ bias for inDrops and mostly uniform with a small 3’ bias for sc-rDSeq. Each gene is divided into 40 segments and counts failing into each segment are calculated for coverage. Ribbon area shows the SEM around mean. **c)** The boxplot displays nUMI/cell for sc-rDSeq and inDrops. On average, sc-rDSeq captures 90k nUMI/cell, inDrops captures 7.4k nUMI/cell. **d)** Distribution of uniquely mapped reads in each type of genomic locations with merged single cells data. **e)** Percentage of genes detected by sc-rDSeq compared to inDrops. **f)** Histogram showing normalized distribution of the fraction of transcripts per biotype expressed in single cells, comparing sc-rDSeq to inDrops. **g)** Cost comparison of sc-rDSeq and other single RNA-seq technologies. Top bars (red) for instrument cost, and bottom bars (blue) for library and sequencing cost per cell. **h)** UMAP of PC9 cells (n = 680) based on total transcriptome, revealing four clusters of populations. **i)** Differentially expressed genes were identified using KNN unsupervised clustering of total transcriptome expression. The average expression of selected marker genes for each cluster are presented in dot plot. **j)** Top five GO terms enriched in each cluster, ranked by fold enrichment. **k)** The average expression of polyA-RNAs showed heterogeneity, including histone RNAs, lncRNAs, sncRNAs, and microRNAs.

Species-mixing experiment with mouse embryonic stem cells (mESCs) and human PC9 cells confirmed that the mixing rate is on the same level as inDrops which is around 5% (**Fig. S2f**). sc-rDSeq showed superior performance in comparison to conventional polyT-based scRNAseq method and is comparable to other state-of-the-art single cell full-length RNA methods, such as VASA-seq and Smart-seq-total, based on the benchmarking results reported previously. We profiled the lung cancer-derived PC9 cells by sc-rDSeq and inDrops to evaluate their technical performance. sc-rDSeq demonstrated higher barcoding efficiency compared to inDrops, with 81% of reads exhibiting the correct library structure. The final mapping rate of reads with proper barcodes, following adapter and rRNA trimming, is 72%, comparable to VASA-drop (∼70%) and higher than Smart-seq-total (∼50%)^10^ (**Fig. S2g,h**). Data from all single cells were combined into pseudo-bulk to assess average gene body coverage across all genes. We observed a pronounced 3’ end bias with inDrops, while sc-rDSeq showed a relatively uniform distribution across the gene body, with a modest 3’ bias due to the polyT5C8N primers (**Fig. 1b**). A similar bias is observed in Smart-seq3 but is less pronounced in Smart-seq-total^11^. Notably, reads from rDS primers alone were evenly distributed (**Fig. S2c**).

Due to the multiple annealing events enabled by sc-rDSeq, on average, sc-rDSeq yielded 90k nUMIs per PC9 cell, representing more than a tenfold increase compared to the 7.4k nUMIs per cell obtained with inDrops (**Fig. 1c**). Gene body coverage analysis showed that sc-rDSeq captured a higher percentage of reads from coding sequences (CDS) and introns, whereas inDrops predominantly captured untranslated regions (UTRs) at the 3’ termini (**Fig. 1d**). Although 75.4% of genes were shared between both methods, sc-rDSeq identified 21.5% unique genes not captured by inDrops, compared to only 3.1% unique genes identified by inDrops (**Fig. 1e**). These unique genes identified by sc-rDSeq were primarily long non-coding RNAs (lncRNAs) and non-polyA RNAs, such as microRNAs (miRNAs) and small non-coding RNAs (sncRNAs), which were largely absent in inDrops **(Fig. 1f; Fig. S2d**).

Despite its improved full-length coverage and total RNA capturing capabilities, the library preparation cost and instrument cost for sc-rDSeq are the same as inDrops. However, the sequencing cost depends on the required coverage, which is approximately doubled due to the method’s higher sensitivity in RNA detection. The estimated costs of sc-rDSeq are $0.08 per cell for library preparation and $0.30 per cell for sequencing (**Supplementary file 1**). The library preparation cost per cell for sc-rDSeq and is six times lower than 10x Chromium ($0.5)^18,19^ and Smart-seq3 (between $0.57 and $1.14)^7^. Since Smart-seq-total utilizes a similar library preparation protocol as Smart-seq3 with additional steps, its costs are expected to exceed those of Smart-seq3. VASA-seq has a slightly higher cost the sc-rDSeq for library preparation ($0.11), but the instrument cost is three times higher than sc-rDSeq due to its two-step microfluidic encapsulation deices. Consequently, sc-rDSeq is the most cost-effective method for single-cell full-length total RNA sequencing so far (**Fig. 1g; Fig. S2e**).

### sc-rDSeq reveals heterogeneous cell populations in cancer cell line by analyzing both polyA and non-polyA RNA

We further investigated the heterogeneity revealed by polyA and non-polyA RNAs using sc-rDSeq. Unique reads were aligned in four sequential rounds, with only unaligned reads passed to the next round: first to small non-coding RNAs, followed by exons, introns, and finally pseudogenes. The expression matrices from each round were then combined for downstream analysis. Applying sc-rDSeq to the NSCLC cell line PC9 identified four distinct cellular states that inDrops could not detect (**Fig. 1h-k**; **Fig. S2i-j**). Clusters 1 and 3 were characterized by more aggressive cancer-related features, such as increased lipid synthesis and cell migration. Cluster 2 displayed heightened activity in the cell cycle, while Cluster 0 exhibited the opposite, with overall lower coverage, indicating a slower metabolic state (**Fig. S2i**). To be more specific, Cluster 0 was enriched for processes like heterochromatin assembly and negative epigenetic regulation of gene expression. Key genes been upregulated like *WAPL*, *SMCHD1* and *BAZ2A*, which promote heterochromatin formation^20–23^ . Cluster 1 showed enrichment for lipid synthesis and proteolysis. Proteases like *CST3*, *CST4*, and *CTSH*, promote tumor aggressiveness^24–26^, while lipid synthesis genes such as *FASN*, *SCD*, and *IDH1* support tumor growth and energy supply ^28–30^. Cluster 2 was enriched for chromosome segregation and proliferation. Highly expressed genes, including *TOP2A*, *AURKA*, and *PTTG1*, were identified as regulators of chromosome instability and cancer progression^27–29^. Notably, various histone RNAs were highly expressed in Cluster 2 (**Fig. 1k**), further supporting roles related to chromosome segregation and rapid cell proliferation. Cluster 3 was linked to cell migration and motility, with genes like *EFNB2*, *ANXA1*, and *LAMC2* promoting migration, invasion, and epithelial-to-mesenchymal (EMT) transition^30–32^. In addition to histone RNAs, other non-polyA RNAs were also differentially expressed across clusters (**Fig. 1f-k**). Cancer-related lncRNAs such as *PVT1*, *GAS5*, and *EBLN3P*^33–36^ was identified. Antisense RNAs, including *HOXA-AS2, TMPO-AS1*, and *ZFAS1*, may regulate their sense RNA counterparts^37–39^.

### DTP cells exhibit G1 arrest and activate different pathways for drug resistance

Previous studies have showed the existences of a small subpopulation of cells, termed "drug-tolerant persisters" (DTPs), that survive short-term exposure to high-dose drug treatment^44^. Recent studies have highlighted distinct fates of DTP cells, but these investigations relied on the conventional scRNA-seq, which only captures polyadenylated RNA and with limited sensitivity^16^. Given the sensitivity of sc-rDSeq in detecting subtle cancer states, we aimed to investigate the emergence of DTPs. As PC9 cells are derived from NSCLC, we focused on two EGFR tyrosine kinase inhibitors (TKIs), Gefitinib and Osimertinib. Gefitinib, a first-generation TKI, targets EGFR mutations but often faces resistance due to secondary mutations like T790M. Osimertinib, a third-generation TKI, overcomes this by targeting both activating and T790M mutations. PC9 cells, which carry an EGFR exon 19 deletion mutation, are highly sensitive to both drugs at low doses. To model drug resistance, we treated PC9 cells with Gefitinib or Osimertinib for nine days at a drug concentration 10-fold greater than the IC_50_ values (**Fig. S3a,b**), eliminating most of the cell population and leaving only a rare subset of DTP cells (<1% of the population). As a control, DMSO was used at equivalent concentrations to mimic treatment conditions without selective pressure.

Using sc-rDSeq, we profiled 7,051 cells across all groups, including 2,982 Gefitinib DTPs, 1,320 Osimertinib DTPs, and 2,749 control cells (**Fig. 2a**; **Fig. S3c,d**). Integration analysis revealed major effects of drug resistance on cell states. The first major finding was that G1 arresting of DTP cells determined by the expression of both classical cell cycle gene and non-polyadenylated histone mRNAs (**Fig. 2a-c)**. Consistent with previous studies^44^, DTPs were mostly in G1 (fraction of G1 cells: Gefitinib 83%, Osimertinib 95.1%), while control cells were largely in G2/M and S phases (fraction of G1 cells: DMSO 4.77%; **Fig. S4b**). Due to the total RNA detection ability, we were able to capture the replication-dependent histone mRNAs which are the only known cellular non-polyadenylated mRNAs. Their expression was elevated in S and G2/M phases (**Fig. 2b**), consistent with histone protein accumulation before mitosis as preparation for division^40^. Cells were then categorized into S and non-S phases based on histone RNA expression levels (**Fig. S4a**). Cell cycle annotation based on cell-cycle and non-polyA histone genes was consistent across both UMAP and population averages expression of histone RNA (**Fig. 2a,c**).

**Fig. 2.**
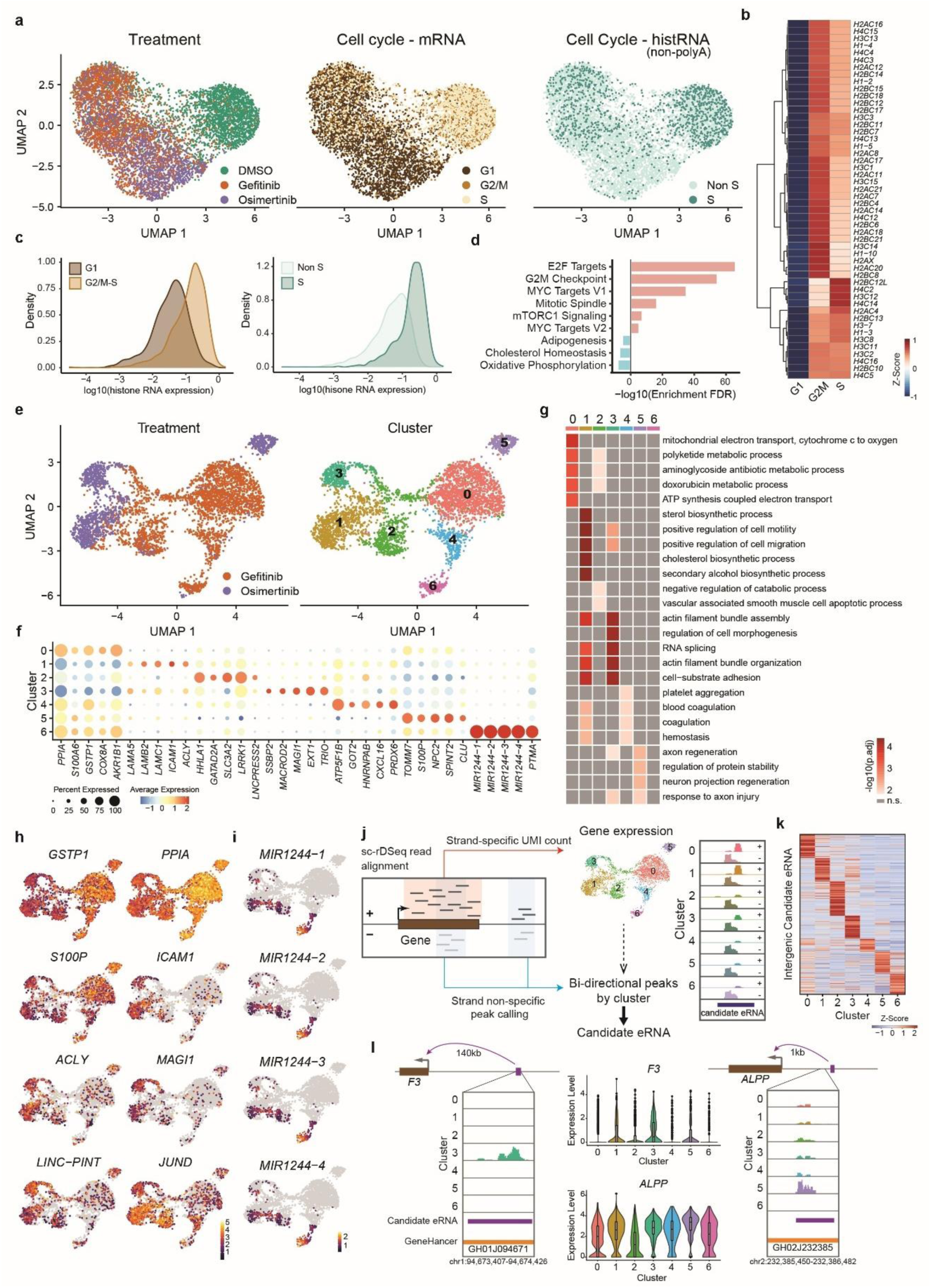
sc-rDSeq reveals heterogeneity based on total RNA in Drug Persistent Cells. **a)** Integrated gene expression profiles of total RNA for PC9 cells under different treatment in UMAP representation colored by treatment (DMSO, Gefitinib and Osimertinib; left), cell cycle affiliation defined by cell cycle marker gene expression (middle), and cell cycle affiliation defined by non-polyA replication-dependent histone mRNA expression (n = 7,051). **b)** Heatmap of histone mRNA expression level (scaled by column) for cells in G1, G2/M or S phases defined by cell cycle marker gene expression. **c)** Distribution of the average expression level of histone mRNA per single cell in G1 and G2M/S phases defined by cell cycle marker gene expression (left) and in S and Non-S phase defined by histone mRNA expression (right). **d)** Significantly up and down regulated MSigDB hallmark (H) gene sets in untreated cells (DMSO) (Enrichment FDR< 0.01). **e)** Subclustering of Gefitinib- and Osimertinib-DTPS. UMAP representation colored by treatment (Gefitinib and Osimertinib; left) and Seurat defined clusters (right) (n = 4490). **f)** The average expression of selected marker genes for each cluster. **g)** Heatmap of significantly upregulated GO BP pathways per cluster, excluding genes for ribosomal proteins. **h)** Example of known marker genes for EGFR TKI resistance or cancer regulation. **i)** Precursor miRNA mir-1244 family are found co-expressed in cluster 6. **j)** Computational pipeline for profiling eRNA expression using sc-rDSeq data. **k)** Heatmap of intergenic candidate eRNA expression in pseudo-bulk clusters aligned with clusters defined by gene expression. Expression levels are normalized by read depth per cluster and then centered by row around zero, displayed as z-scores. Rows are ordered based on agglomerative clustering results. **l)** Example of differentially expressed eRNAs at the pseudo-bulk level, where the targeted gene is also differentially expressed at the single-cell level.

To identify cellular expression programs associated with drug resistance, we analyzed differentially expressed gene ontology pathways between DTPs and control cells (**Fig. 2d**). Six proliferation-related pathways were upregulated in control cells (MYC targets v1, G2M checkpoints, E2F targets, MYC targets v2, and Mitotic spindle), consistent with their role in cancer drug responses^41^. We further investigated the heterogeneity within DTPs through subclustering treated cells. UMAP analysis revealed seven distinct clusters, with Gefitinib DTPs exhibiting greater cellular diversity, predominantly occupying clusters 0, 2, 4, and 6. Osimertinib DTPs were primarily concentrated in clusters 3 and 5, while cluster 1 comprised cells from both treatments, with a higher proportion of Gefitinib-treated cells (**Fig. 2e**; **Fig. S4c-e**). The differentially expressed genes enriched for different biological processes that suggests several strategies for cells to bypass the drug treatment. Clusters 0, 5, and 6 displayed elevated metabolic pathway expression, including cytoplasmic translation and ribosome biogenesis, indicating increased protein synthesis requirements for the development of resistance^42,43^ (**Fig. 2g**; **Fig. S4h**). These clusters also showed higher MYC target expression and oxidative phosphorylation (**Fig. S4f,g**), consistent with previous findings in Gefitinib-resistant NSCLC cells^49^. Notably, several genes upregulated in these clusters, such as *PPIA*, *S100A6*, and *GSTP1* (**Fig. 2f,h**), have been previously linked to EGFR TKI resistance in cancer^44–46^. Cluster 5 showed higher expression of tumor suppressor genes, including *S100P*, *NPC2*, *SPINT2*, and *CLU*^47–50^ (**Fig. 2f,h**), suggesting reduced cellular aggressiveness. While cluster 6 shares most differentially expressed genes with cluster 5, it uniquely expressed four precursor microRNAs (*MIR1244*-1/2/3/4) that form mature mir-1244 (**Fig. 2i**). mir-1244 is involved in mRNA degradation and translation inhibition, which has previously been reported to sensitize cells to cisplatin, induce resistance, and cause cellular senescence in multiple cancers^51–53^. Clusters 1 and 3 exhibited upregulation of tumor-aggressive pathways, including EMT transition, cholesterol homeostasis, and extracellular matrix (ECM) remodeling^54^ (**Fig. 2g**; **Fig. S4f-h**). Genes been upregulated include *ACLY* for lipid metabolism^55^, laminin subunits *LAMA5*, *LAMB2*, *LAMC1* and the cell surface glycoprotein gene *ICAM1* for ECM formation (**Fig. 2f,h**). Cluster 3 also present tumor suppressive activities with the expression of tumor suppressors like *LINC-PINT*, *MAGI1*, *SSBP2*, *MACROD2*, and *EXT1*^56–60^. These findings suggest an intriguing phenomenon in which the persistent cells of cluster 3 exhibit both tumor suppressor and aggressive activities. Interestingly *JUND*, part of the AP-1 transcription factor family, was expressed in both aggressive (clusters 1 and 3) and less aggressive (clusters 2 and 4) states (**Fig. 2h**). This highlights its dual role in promoting or inhibiting growth depending on the cellular context^61^.

Finally, to investigate whether chromosomal copy number variations (CNVs) act as genomic drivers of resistance, we analyzed single DTP cells using inferCNV^62^ with DMSO-treated cells as a reference. Most Gefitinib- and Osimertinib-DTP cells exhibited chr19 gain and partial loss of chr2, except for two rare subpopulations, cluster 2 and 3 (**Fig. S3e**). Cells from CNV cluster 2 and 3 were primarily concentrated in cluster 3 and cluster 1 on the gene expression UMAP, respectively. Other CNV-based clusters showed some tendency to be restricted to specific transcriptional clusters but did not display a clear correlation with gene expression (**Fig. S3e**). Overall, subpopulations identified through RNA expression exhibited greater variation than those inferred from CNV analysis. It aligns with previous studies^63^, suggesting that CNVs are just one of the mechanisms cancer cells use to develop drug resistance, and a deeper understanding of the underlying expression-based and mutational pathways is instrumental. These findings highlight the potential of sc-rDSeq in identifying drug-resistant cells and optimizing combination therapies that target cluster-specific pathways.

### Enhancer RNAs align with gene expression variability

Enhancers, distal regulatory elements often disrupted in cancer, produce short noncoding RNAs called enhancer RNAs (eRNAs) that influence gene expression^64,65^. However, detecting eRNAs at the single-cell level is difficult due to their short length, low abundance, and instability. Recent advances like scGRO-seq have improved detection but remain limited in throughput^66^. Given the strand-specificity of sc-rDSeq, we explored its potential for profiling eRNAs by identifying bidirectionally transcribed regions. Since the eRNAs are cell-type-specific and expressed in low amount, we extracted bidirectional peaks from aggregated reads per cluster (2%-4.4% bidirectional), merging them across clusters to compile a candidate eRNA list (**Fig. 2j**; **Fig. S5a**). In DTP cells, we identified 9,750 candidate eRNAs, with 6% in intergenic regions and 25–30% previously annotated^67^ (**Fig. S5b,d**), mostly located near transcription start sites (TSSs) and extended up to 50 kb upstream or downstream of protein-coding genes and lncRNAs (**Fig. S5c,e-f**). We then quantified eRNA expression at the pseudo-bulk level. While intergenic eRNAs were not inherently cluster-specific, their expression patterns correlated well with gene expression clustering (**Fig. 2k**; **Fig. S5h**). Based on GeneHancer^67^, an experimentally validated regulatory database, we linked candidate eRNAs to their putative target genes and found significant overlaps with differentially expressed genes (**Fig. S5g**). For instance, a cluster 3-specific eRNA targeted *F3* (coagulation factor III), which upregulated in the same cluster. Another cluster5- specific eRNA near *ALPP* (1 kb away), which showed highest expression in cluster 5, consistent with its role in Gefitinib response^68^ (**Fig. 2l**).

### Extraction of Alternative Splicing Events to Dissect DTP Subpopulations

Alternative splicing (AS) drives phenotypic diversity and cancer progression by generating multiple transcripts from a single gene^69–71^. Detecting AS at the single-cell level has been challenging due to limited transcript coverage in 3’ end methods and low sensitivity in full-length sequencing^72,73^. Leveraging the full-length coverage provided by sc-rDSeq, we evaluated its capacity to quantify AS events. sc-rDSeq detected four times more splicing events than inDrops, primarily skipped exons, while capturing most inDrops-identified events (**Fig. S6a,b**). To assess cell-state-dependent splicing in DTP cells, pseudo-bulk data from gene expression clusters were analyzed with RSEM^74^, revealing significant isoform switching in 20–38% of expressed genes per cluster (**Fig. 3c**). For example, while total *MDK* expression was comparable between cluster 3 and 4, cluster 4 predominantly expressed full-length *MDK-203*, whereas cluster 3 favored *MDK-204*, which skips exon 3 and results in truncated tMDK (**Fig. 3a**). Midkine is a secreted protein that functions as growth factor involved in tumorigenesis^75^. Despite lacking the N-terminal domain, tMDK retains its functional C-terminal domain and was shown to accelerate tumor formation in mice^76,77^. Notably, cluster 3 displays higher EMT levels (**Fig. S4g**), suggesting the unexplored role of tMDK in this process. Similarly, *CTSH*, encoding Cathepsin H, a protease promoting metastasis^26,78^, showed higher expression in cluster 4, which primarily expressed full-length *CTSH-207*. Cluster 2 favored *CTSH-236*, a shorter isoform lacking the inhibitory propeptide domain, implying enhanced protease activity. Single-cell isoform analysis remains challenging due to low read depth, but doing PCA analysis on isoform-switched genes largely recapitulated gene expression-based clustering (**Fig. 3d,e**; **Fig. S6c-f**). Intriguingly, cluster 0 split into two subclusters in isoform UMAP, revealing AS-driven heterogeneity. Single-cell isoform expression validated pseudo-bulk trends for *MDK* and *CTSH* (**Fig. 3f,g)**. Notably, the longer isoform CTSH-207 had very low expression on single cell level due to the need of higher sensitivity over 5’ exons, suggesting the need of pseudo-bulk level analysis to better reveal the AS events. Our findings underscore the utility of sc-rDSeq in resolving AS heterogeneity at single-cell resolution. Isoform switching correlates with distinct cell states in DTP populations, with truncated isoforms potentially contributing to functional plasticity during adaptation. While pseudo-bulk analysis enhances detection of low-abundance isoforms, integrating isoform-level data refines transcriptional landscapes, highlighting AS as a critical layer of tumor cell heterogeneity.

**Fig. 3.**
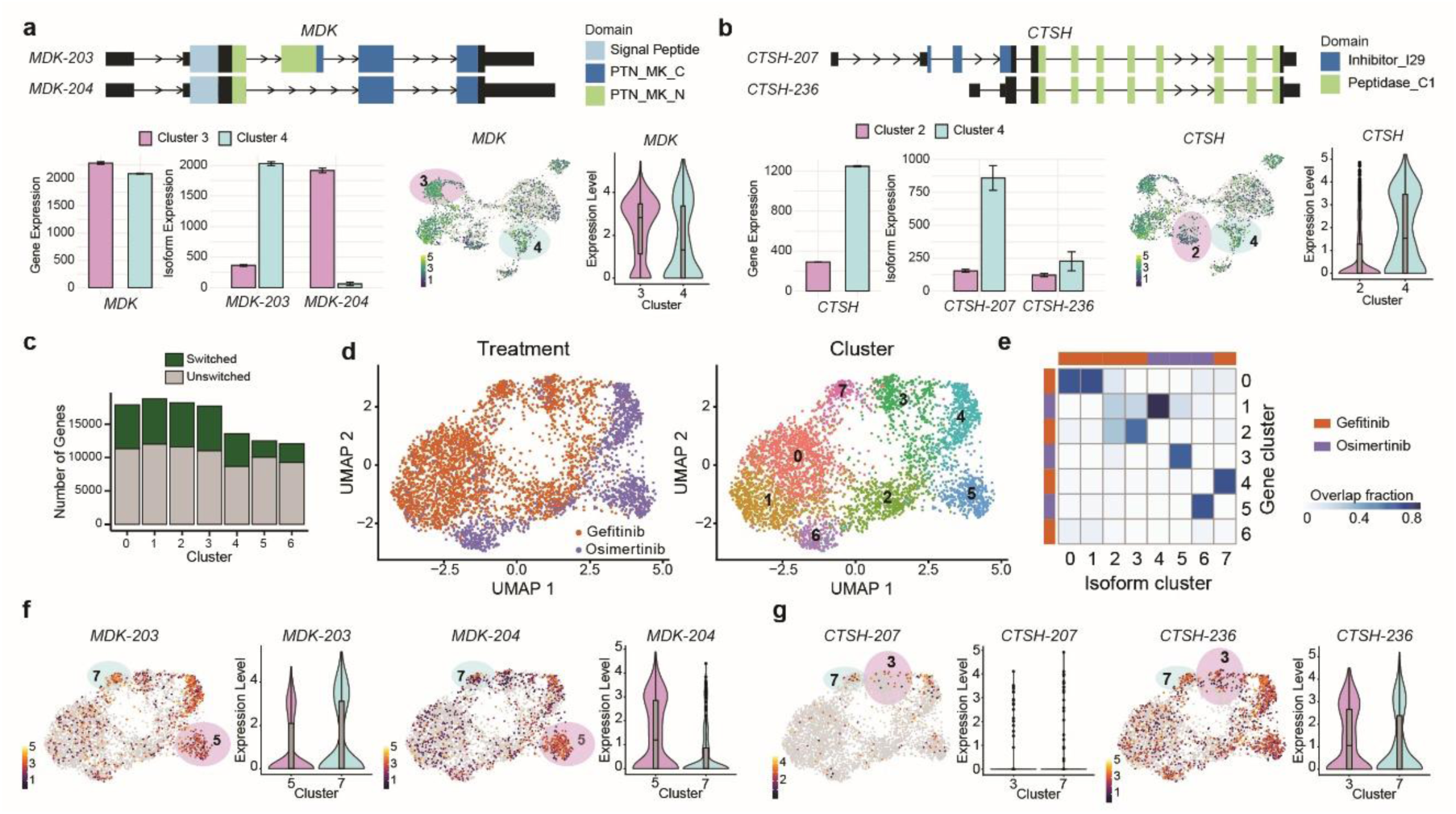
Alternative isoform usage adds another layer of heterogeneity to DTPs cells. **a)** and **b)** Isoform switching in *MDK* (a) and *CTSH* (b) genes, with isoform structures (top), gene expression in pseudo-bulk (bottom left), and single-cell expression in clusters 3 & 4 (*MDK*) or clusters 2 & 4 (*CTSH*) (bottom right). **c)** Number of genes with isoform switching in pseudo-bulk clusters. **d)** UMAP visualization of isoform expression in PC9 cells under different treatments, colored by treatment (left) and Seurat-defined clusters (right) (n = 4,490). **e)** Heatmap showing the overlap fractions between clusters defined by gene and isoform expression, with rows and columns colored by treatment groups. **f)** and **g)** *MDK* (f) and *CTSH* (g) isoform expression (*MDK-203/MDK-204, CTSH-207/CTSH-236*) on the isoform UMAP and violin plots.

### Genetic variation analysis reveals non-driver role in DTP heterogeneity

To evaluate genetic contributions to drug resistance, we analyzed single nucleotide variants (SNVs) and indels in DTPs and control cells using mutect2^79^ on pseudo-bulk data. We start with screening mutations in *EGFR* gene (**Fig. S7a,b**), and found the site for the canonical resistance variant T790M (c.2369C>T) was unmutated both before and after drug treatment, indicating other sources for drug resistance. On the other hand, a recurrent 3’ UTR SNV (chr7:55206391 T>C) dominated across groups, which might have an influence on translation efficiency of EGFR (**Fig. S7a**). Moreover, most EGFR SNVs in untreated cells were absent in DTPs, suggesting these mutations are not contributing to drug sensitivity (**Fig. S7b**). We then move on to look for all SNVs that potentially contribute to the drug resistance. Given the short time of drug treatment (9 days), we don’t anticipate the accumulation of de-novo SNVs in the DTPs. Accordingly, we focused on the pre-existing SNVs and indels with altered allele frequencies post-treatment. To identify mutations that may have contributed to drug resistance but were pre-existing in the untreated (DMSO) state, we used the human genome assembly GRCh38 as the reference. We identified 857,033, 29,851 and 22,474 SNVs and indels in DMSO, Gefitinib and Osimertinib DTPs, respectively. The SNVs are predominantly found in introns and 3’UTR (**Fig. S7c**). After filtering low-expression variants and significancy testing, we identified 59 enriched and 41 depleted SNVs in Gefitinib DTPs, and 24 enriched and 21 depleted in Osimertinib DTPs, with 15 enriched and 4 depleted SNVs in both treatment groups (**Fig. 4a; Fig. S7d**). Although the 3’ UTR mutations could have significant impact in gene regulation, we focused on the 17 missense and splice-site mutations to reveal the direct functional relevance (**Fig. 4b**). We found that many (8/14) of these mutations were found in genes related to lung cancer^80–85^. For example, knockdown *MUC16* is shown to activate the downstream targets of EGFR pathway^86^, the enriched allele frequency in Gefitinib DTPs of *MUC16* is observed and potentially reflecting compensatory adaptation to EGFR inhibition. Conversely, *IGFBP6* mutation allele frequency was depleted, suggesting selective pressure against growth factor loss (**Fig. 4c**). We then performed single cell genotyping over these SNVs using SComatic^87^ (**Fig. 4d; Fig. S7e**). However, it showed no clear correlation with transcriptional states, implying that SNVs are not the major drivers for transcriptional heterogeneity but rather a necessity to achieve resistance. Together, these findings, although still limited in coverage, highlight the limited role of genetic variation in shaping DTP cell states and underscore non-mutational mechanisms of resistance.

**Fig. 4.**
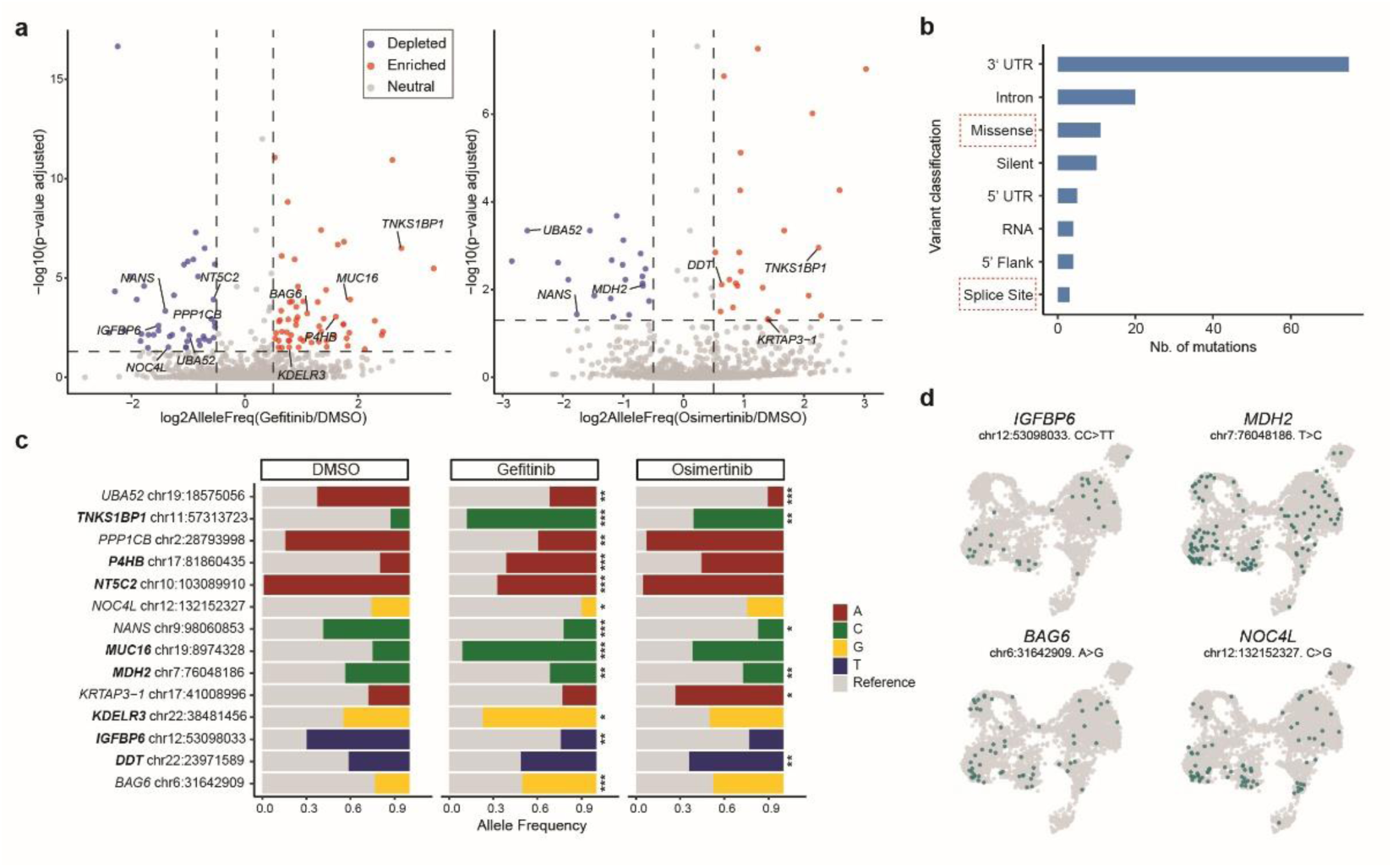
Genetic variation analysis reveals non-driver role in DTP heterogeneity. **a)** Volcano plot showing fold changes of mutation allele frequency comparing Gefitinib (left) and Osimertinib (right) DTPs to DMSO. Log2FC threshold is 0.5 and adjusted p-value threshold is 0.05. Missense and splice site mutations are highlighted. **b)** Variant classification counts of enriched and depleted mutations identified in (a). **c)** Allele frequency of selected SNVs in three groups, with genes related to lung cancer highlighted. **d)** UMAP visualization highlighting cells carrying selected SNVs.

## DISCUSSION

sc-rDSeq is a ground-breaking droplet-based, ribosomal-depleted sequencing method designed to capture a wide spectrum of single-cell transcriptomes with high sensitivity and exceptional cost-effectiveness. It provides full-length coverage of total RNA, including both polyA+ and polyA- transcripts, which are often overlooked by traditional scRNA-seq methods. sc-rDSeq matches state-of-the-art methods like Smart-seq-total and VASA-seq in performance with improved scalability, accessibility, and cost-efficiency. Compared to the gold-standard polyT-based method inDrops, sc-rDSeq significantly improves full-length coverage, diverse RNA profiling, and sensitivity, achieving a tenfold increase in UMIs. sc-rDSeq does not require sophisticated devices beyond basic microfluidic setups used in standard droplet-based scRNA-seq, making it highly compatible with most commercially available platforms while overcoming the lower encapsulation efficiency seen in VASA-drops due to the droplet merging step. By excluding rRNA transcripts at intial capture step, sc-rDSeq eliminates the need for dedicated rRNA depletion, reducing costs and enhancing total RNA amplification efficiency. The high throughput of sc-rDSeq enabled us to investigate the emergence of drug persistence in cancer cells, revealing a complex landscape of cellular responses to EGFR tyrosine kinase inhibitors. Specifically, we confirmed G1 cell cycle arrest in DTPs not only through conventional cell cycle markers but also via polyA- histone mRNAs. Additionally, we identified rare cell populations marked by non-coding RNA like microRNAs, and demonstrated that the expression of intergenic eRNAs is aligned with gene expression-based clusters. The full gene-body coverage facilitated the extraction of AS events, linking transcript diversity to functional protein potential. Furthermore, sc-rDSeq expanded the detection power of SNVs, allowing the identification of mutations with allelic fractions enriched upon drug treatments. A current limitation of sc-rDSeq is its restriction to human cells due to the design of the rDS primers. Expanding its application to other species requires only redesigning barcoded beads with rDS specific to the target species. Additionally, improving sensitivity will enhance the accurate detection of alternative splicing, eRNAs, and SNVs at the single-cell level. To address this, we plan to integrate sc-rDSeq with CloneSeq^88,89^ in future studies. Profiling 3D-cultured clones of untreated cells versus DTPs originating from individual cells should enhance sensitivity, improve coverage, and reduce signal-to-noise ratios by minimizing biological variability, such as cell cycle effects and transcriptional bursting. sc-rDSeq, particularly when integrated with CloneSeq, will significantly enhance our ability to dissect cancer biology and serve as a powerful tool for personalized cancer medicine and research. As we continue to refine sc-rDSeq and explore its applications, we anticipate it will become an indispensable tool in the field of single-cell sequencing.

## ACKNOWLEDGMENTS

O.R. is supported by research grants from the European Research Council (ERC, # 715260 SC-EpiCode), the Israeli Center of Research Excellence (I-CORE) program, the Israel Science Foundation (ISF, #1618/16), and Azriely Foundation Scholar Program for Distinguished Junior Faculty.

## AUTHOR CONTRIBUTIONS

X.S. and O.R. conceived the study, prepared the figures and wrote the manuscript. D.E. and M.A. conducted the prove-of-concept of the experiments. X.S., and O.R designed the experiments.

X.S. performed tissue culture, drug resistance experiments and single cell library preparations.

X.S. and W.F. performed microfluidics operation and sequencing. X.S. conducted the benchmarking, gene expression, eRNA and isoform analysis. S.D. conducted the somatic mutation analysis. E.R. conducted the copy number variation analysis. A.A. and D.E. reviewed the manuscript.

## RESOURCE AVAILABILITY

### Lead contact

Further information and requests for resources and reagents should be directed to and will be fulfilled by the lead contact, O.R. and X.S.

### Materials availability

This study did not generate new unique reagents.

### Data and code availability

The accession number for the data reported in this paper is GEO: GSE282074.

## METHODS

### Cell culture

Human PC9 lung adenocarcinoma cells were kindly provided by Prof. Ravid Straussman (Weizmann Institute, Israel). PC9 cells were grown in DMEM (Sigma-Aldrich, cat #D5671) supplemented with 10% fetal bovine serum (FBS; Biological Industries Israel, cat #04-007-1A), 50 μg/ml penicillin-streptomycin (Biological Industries Israel, cat #03-031-1B), 2 mM L-glutamine (Biological Industries Israel, cat #03-020-1B), and 1 mM sodium pyruvate (Biological Industries Israel, cat #03-042-1B). Mouse embryonic stem cell R1 were kindly provided by Prof. Eran Meshorer (The Hebrew University, Jerusalem). R1 cells were grown on 0.1% gelatin-coated standard tissue culture dishes and maintained in ESC medium (DMEM, 15% ESC-grade FBS, 50 μg/ml penicillin-streptomycin, 2 mM L-glutamine, 1 mM sodium pyruvate, 0.1 mM non-essential amino acids (Biological Industries Israel, cat #01-340-1B), 0.1 mM β-mercaptoethanol (Sigma-Aldrich, cat #M3148))

### Viable cell counting by colorimetric assay and dose-response curve

The dose-response was assessed by counting viable cells using Cell Proliferation Kit (XTT based; Biological Industries Israel, cat #20-300). Cells were seeded in 96-well plates at a concentration of 5,000 cells/well. 24 hours after plating, media was removed and replaced with media containing Osimertinib (AZD9291, Selleck Chemicals, cat #S7297) or Gefitinib (Selleck Chemicals, cat #S1025) for 72 hours. Both drugs were tested at concentrations ranging from 1 nM to 100 μM, using 3-fold serial dilutions. Absorbance at 450 nm (sample absorbance) and 660 nm (reference absorbance) was measured in two steps using a spectrophotometer (Biotek Synergy H1 plate reader). A blank control, containing only medium, was set up for each row. The effective absorbance of each well was calculated by: 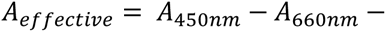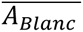, and final viability is calculated by : 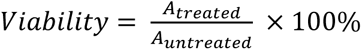. The drug concentration required to inhibit cell growth by 50% (IC50) was determined from concentration-response curves generated with the R package drc v3.0-1^90^.

### Drug treatment experiments

EGFR inhibitors were prepared as 10 mM stock solutions in DMSO: Osimertinib and Gefitinib. Small aliquots (10-15 µL) were stored at -20 °C to minimize freeze-thaw cycles. For drug treatment experiments, stock solutions were first diluted in DMSO and then in culture medium, maintaining a final DMSO concentration of 1% across all dilutions. Untreated cells were given 1% DMSO as a control. As start, 1 x 10^6^ PC9 cells were seeded to a 10 cm^2^ plate. 24h after seeding, 1 μM Osimertinib and 1 μM Gefitinib were given to cells. Every 3 days, medium was changed with fresh drugs. After 9 days, remained cells were collected for sc-rDSeq. Control groups were treated with DMSO for 3 days.

### Species-mixing experiments

To determine off-species contamination in the single-cell preparations, sc-rDSeq was performed with a suspension mixture of PC9/R1 mESCs cells. The suspension mixtures were prepared at a total concentration of 300,000 cells/ml (1:1 ratio).

### Microfluidics device design, fabrication and operation

Two microfluidics devices were used: (I) to produce acrylamide hydrogel microparticles with acrydite-modified DNA primers for barcoding; (II) to encapsulate single cells with barcodes, lysis buffer, and reverse transcriptase (RT) enzyme within droplet. Device design, fabrication and operation protocols followed the protocol described previously^88,89^.

### Production of sc-rDSeq barcoded hydrogel beads and quality control

Hydrogel beads carrying barcoded DNA primers were produced following the method described before^88,89^. The acrydite-modified DNA primer and the hydrogel beads containing it were handled in darkness or under UV-filtered light. DNA primers on polymerized hydrogel beads were barcoded using a combination of the split-and-pool method and a primer extension reaction. This process involved three rounds of priming: the first two rounds assigned single-cell barcodes, and the third round extended the rDS/polyTN8 primers. Each round consisted of primer loading, hybridization and extension, performed by the Biomek 4000 liquid handling robot (Beckman Coulter Life Sciences).

In the first two rounds, acrydite hydrogels with 2.5×isothermal amplification buffer (NEB, cat #B0537) and 0.85 mM dNTP (NEB, cat #N0446) were distributed in a 384-well plate at 6 µL per well (∼40,000 hydrogel beads). For each round, 9 µL of 50 µM BC1 primers (PE1*-BC1*-Spacer1*, detailed sequences at Table 1) or BC2 primers (Spacer1*-BC2*-UMI*-Spacer2*, detailed sequences at Table 1) were added to each well, with each well containing a unique primer. After denaturation at 85°C for 2 minutes and hybridization at 60°C for 20 minutes, 5 µL of primer extension mix containing 1.8 U of DNA Polymerase (NEB, cat #M0537) and 0.3 mM dNTP (NEB, #N0446) in 1× isothermal amplification buffer (NEB, cat #B0537) was added, followed by incubation at 60°C for 60 minutes. The reaction was stopped by adding 20 µL of stop buffer (100 mM KCl, 10 mM Tris-HCl [pH 8.0], 50 mM EDTA, 0.1% [v/v] Tween-20) and incubating at room temperature for 10 minutes. The beads were then collected and washed for single strand as described previously (Klein et al., 2015). At each round, before washing, 0.5 µL of beads were saved in 9.5 µL DDW in a 1.5 mL tube for later-on quality control. The split-and-pool process produced a barcode combination complexity of 147,456 (Supplementary Figure S1 A, C). In the third round, a primer mixture was prepared for all beads, consisting of 50 µM rDS/polyTN8 primers (Spacer2*-UMI*-rDS* primer/Spacer2*-UMI*-polyTN8* primer, detailed sequences at Table 1) in a 4:1 ratio. The rDS primers were pre-mixed from 220 different sequences (Supplementary File 2.sc-rDSeq Barcode primers). The priming procedure was like the first two rounds, with the exception that, each well contained the same primer mixture (Supplementary Figure S1 A-C).

**Table 1.**
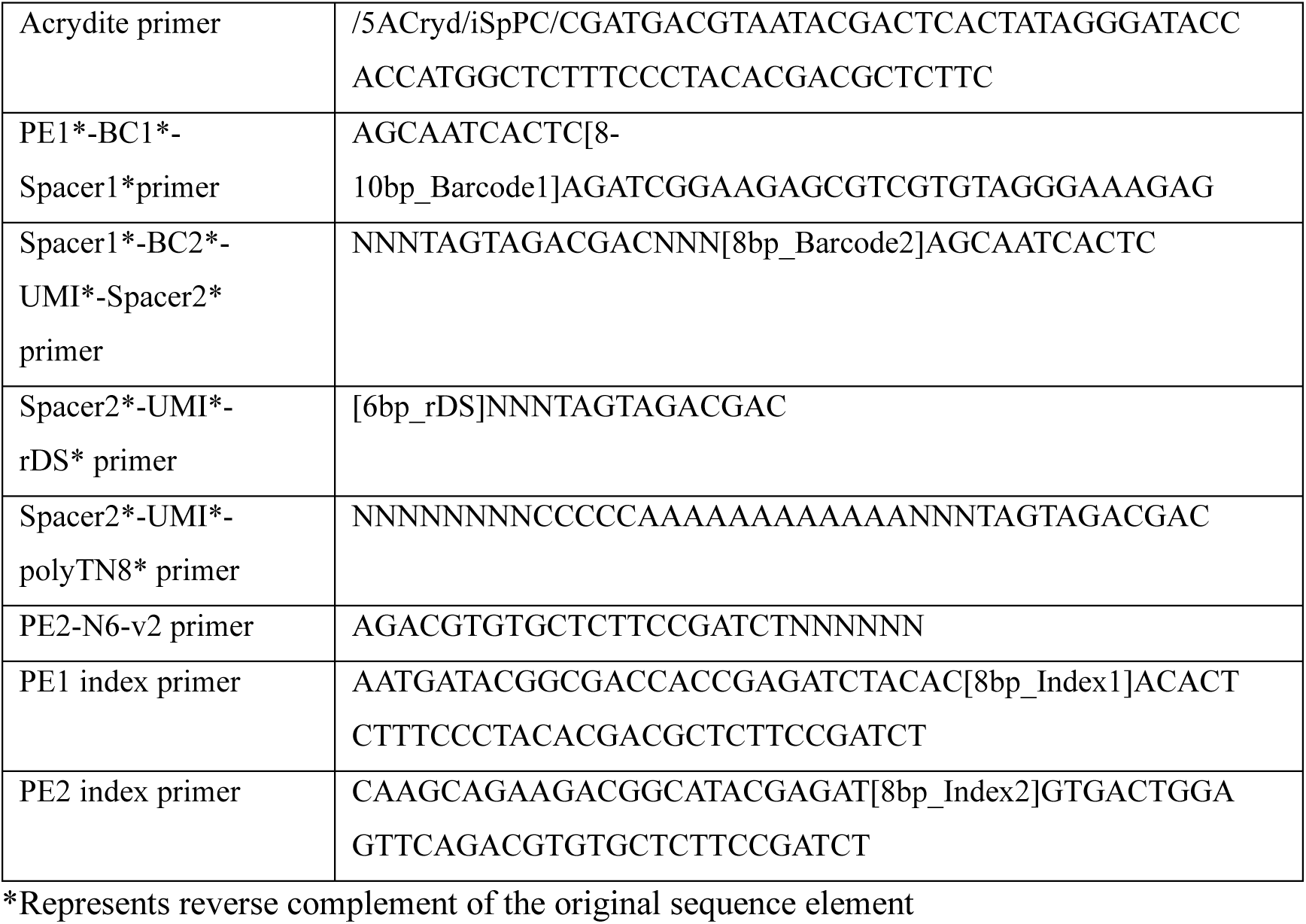
Primer list.

To remove primers that failed to elongate at each step and did not end with rDS/polyTN8, a cleanup process was performed. After priming, all beads were collected in a 50 mL tube and washed twice in T10E0.1T0.1 buffer (10 mM Tris-HCl [pH 8.0], 0.1 mM EDTA, 0.1% [v/v] Tween-20). Next, 0.25 U/µL ExoI and 1×ExoI buffer (NEB, cat #M0293) were added. The tube was rotated at 37°C for 60 minutes in the dark. The reaction was stopped by adding 10 mL stop buffer and incubating on ice for 10 minutes, followed by washing as in the previous step.

After barcode synthesis, the barcoded hydrogel beads were filtered twice using a 70 µm cell strainer (pluriSelect, cat #43-10070-50) to obtain homogeneously sized beads with an approximate diameter of 60 µm and an average of 10^9^ copies of fully extended DNA primers per bead^91^. Small batches of beads (∼0.5μl) saved per round were first exposed to UV for 10 minutes to release the primers, then run on a 2% agarose gel for quality control. Clear bands around 120 bp should be observed for the final products, while bands from earlier steps may show slight smears at lower sizes due to incomplete primer extensions.

### Single-cell barcoding

For both inDrops and sc-rDSeq, the microfluidics operations following protocol described previously^88,89^. The devices have four inlets: 1) Cell inlet: cells were loaded at 300,000 cells/ml in PBS containing 10% v/v OptiPrep (STEMCELL Technologies, cat # 07820) and maintained in suspension using a magnetic micro-stirrer bar placed within the tube. 2) Barcoded hydrogel beads inlet: inDrop barcoding beads were prepared as previously described^91^, and sc-rDSeq barcoding beads were prepared in the last step. Around 100-150 µL beads were centrifuged in a 1.5-ml Eppendorf tube at 1500 rcf for 2 min to obtain packed beads. After aspirating residual buffer from the pelleted beads, the tube was loaded onto the corresponding inlet in the microfluidics setup. 3) Reverse transcription/lysis mix inlet: 150µL RT/lysis mix consisted of 30 µL 10X StellarScript buffer, 15µL StellarScript RT enzyme 200 U/µL (StellarScript HT Reverse Transcriptase Kit, Watchmaker cat # WM-7K0070), 10 µL murine RNase inhibitor (NEB, cat #M0314) , 15 µL 1 M Tris-HCl (pH 8.0), 10 µL 0.1 M DTT, 9 µL 10% (v/v) IGEPAL CA-630 (Sigma-Aldrich, cat #I8896), 6 µL 25 mM dNTPs (NEB, cat #N0446S) and 55 µL nuclease-free water. 4) Carrier oil inlet: The carrier oil was 3 ml of HFE-7500 with 2% (w/w) fluorosurfactant. During microfluidics runs, cell suspension and collection tubes were kept on ice. The microfluidics chips generate monodispersed droplets with volumes in the range of 1-3 nL. inDrops collection emulsions were kept in 1.5 mL Eppendorf tubes pre-filled with 200 µL 2% HFE-7500, and sc-rDSeq collection emulsion were collected into 96-well plates pre-filled with 50 µL 2% HFE-7500 per well. After collections, 100 µL and 50 µL mineral oil were added on each collection respectively.

### Library preparation

The inDrops libraries were prepared following the protocol described previously^91^. The sc-rDSeq process was similar, with the following modifications: after the microfluidics collections, the collection well plate was exposed to 6.5 J/cm²from a 365-nm UV lamp while kept on ice for 10 min to release photocleavable barcoding primers from the barcoding beads. Next, the collection well plate containing the UV-exposed emulsion was transferred to a PCR machine for a ramping-cycling reverse transcription reaction under 20 cycles of the following conditions: 8 °C for 12 s, 15 °C for 45 s, 20 °C for 45 s, 30 °C for 30 s, and 50 °C for 15 s. Then, emulsions from 2-3 wells were combined into a 1.5 mL Eppendorf tube, with each tube containing 1,000-3,000 cells per sample. The following protocol, up to and including *in vitro* transcription (IVT), was the same as for inDrop^91^.

After IVT, resulting aRNA was purified using a 1.3X reaction volume of AMPure XP beads and eluted with 20 µL of 10 mM Tris-HCl. An aliquot of 9 µL was stored at -80 °C as a backup. Next, RNA was fragmented using the RNA fragmentation kit (Invitrogen, cat #AM8740). 9 µL of aRNA was combined with 1 µL of RNA fragmentation reagent and incubated at 70 °C for 1min, then transferred to ice. A 40 µL fragmentation stop mix containing 5 µL fragmentation stop solution and 35 µL 10 mM Tris-HCl was added. Fragmented RNA was purified with a 1.3X reaction volume of AMPure XP beads and eluted in 10 µL of 10 mM Tris-HCl. The amplified and fragmented RNA was reverse transcribed using a random hexamer primer. Specifically, 12 µL RNA was mixed with 2 µL of 100 µM PE2-N6-v2 primer containing random hexamer (Table 1), 1 µL of 10 mM dNTP, 2 µL of 10X StellarScript buffer, 1 µL of 0.1 M DTT, 1 µL Murine RNase inhibitor (40 U/µL), and 1 µL of StellarScript RT enzyme (200 U/µL). Samples were incubated at 25 °C for 10 min, 50 °C for 60 min, and 80 °C for 10 min. Following reverse transcription, the reaction volume was increased to 40 µL by adding 20 µL of nuclease-free water. An aliquot of 20 µL was stored at -80 °C for backup. The resulting cDNA was purified with a 1.2X reaction volume of AMPure XP beads and eluted in 11.5 µL of 10 mM Tris-HCl. The libraries were then PCR amplified using a standard mix. Primers containing Illumina library indices for multiplexing were added. Each PCR reaction contained 11.5 µL of the post-reverse transcription cDNA library, 12.5 µL of 2X KAPA HiFi HotStart ReadyMix (Roche), and 1 µL of 25 µM PE1/PE2 index primer mix (Table 1), and consisted of 10 amplification cycles, following conditions: 98°C for 2 min, 2 x (98°C for 20 s, 55°C for 30 s, 72°C for 40 s), 8 x (98°C for 20 s, 65°C for 30 s, 72°C for 40 s), 72°C for 10 min. Amplified libraries were purified using a 0.7X reaction volume of AMPure XP beads and eluted in 20 µL of nuclease-free water. Library quality was confirmed on the Agilent 2200 TapeStation nucleic acid system (Agilent) using the Agilent High Sensitivity D1000 DS DNA kit. The final libraries had an average size of 300-500 bp.

### Library sequencing

Two types of sequencing platforms were used in this work, based on the data size. inDrops libraries were sequenced using Illumina NextSeq 2000 instrument with NextSeq 1000/2000 P2 Reagent kit (100 Cycles). sc-rDSeq were sequenced at Hadassah Genomic center using NovaSeq 6000 instrument with NovaSeq 6000 S1 Reagent Kit (100 cycles). Cycle distribution in both cases was 50, 70, 8, 8 cycles for Read 1, Read 2, index 1 and index 2 respectively.

#### Raw data filtering and demultiplexing

The raw sequencing data were paired-end fastq files, split into two lanes and four lanes per read for the NextSeq 2000 and NovaSeq 6000, respectively. In data QC, Read1 from all lanes was iterated through and filtered for reads possessing the correct barcode structure (for inDrops, search for the W1 adaptor; for sc-rDSeq, search for Spacer1 and Spacer2), and single-cell barcodes and UMIs were extracted. Barcodes that could be uniquely assigned to an accepted barcode with a Hamming distance of 2 nt or less were merged. UMI deduplication was performed by removing reads with duplicated UMIs under the same single-cell barcode. Unique UMI reads per barcode were counted and plotted in a histogram. Thresholds were determined based on the separation between the first and second peaks. This threshold was then applied in the filtering step, where a similar process was performed as in QC, but outputting reads with UMI counts greater than the threshold into barcode-specific fastq files. Single cell fastqs were then trimmed using cutadapt v4.0^92^ for Illumina adaptors, barcode spacers and repeated elements (20x A/T/G/C). Finally reads aligned to ribosomal RNA were removed using bwa mem and bwa aln v0.7.17^93^ with human 12S, 16S, 18S, 28S and 5.8S rRNA sequences from NCBI.

#### Read alignments

inDrops reads were aligned as described previously^88,89^. For sc-rDSeq, reads were aligned in multiple rounds to exclusively map to total RNA. Briefly, reads were first mapped to the whole genome (GRCh38, Ensembl Release 112), then counted with different gene annotation GTF files in each round. Only reads that failed to align proceeded to the next round. The alignment order was short RNA, long RNA exons, long RNA introns, and pseudogenes. GTF files used for each round were subsets of the full GTF, created by defining specific filters for gene biotypes as listed in Table 2. Different alignment tools were applied in each step to maximize alignment efficiency. Alignments were performed in a strand-specific manner. To analyze gene coverage, BAM files from individual cells were combined and qualified by QoRTs^94^.

**Table 2.**
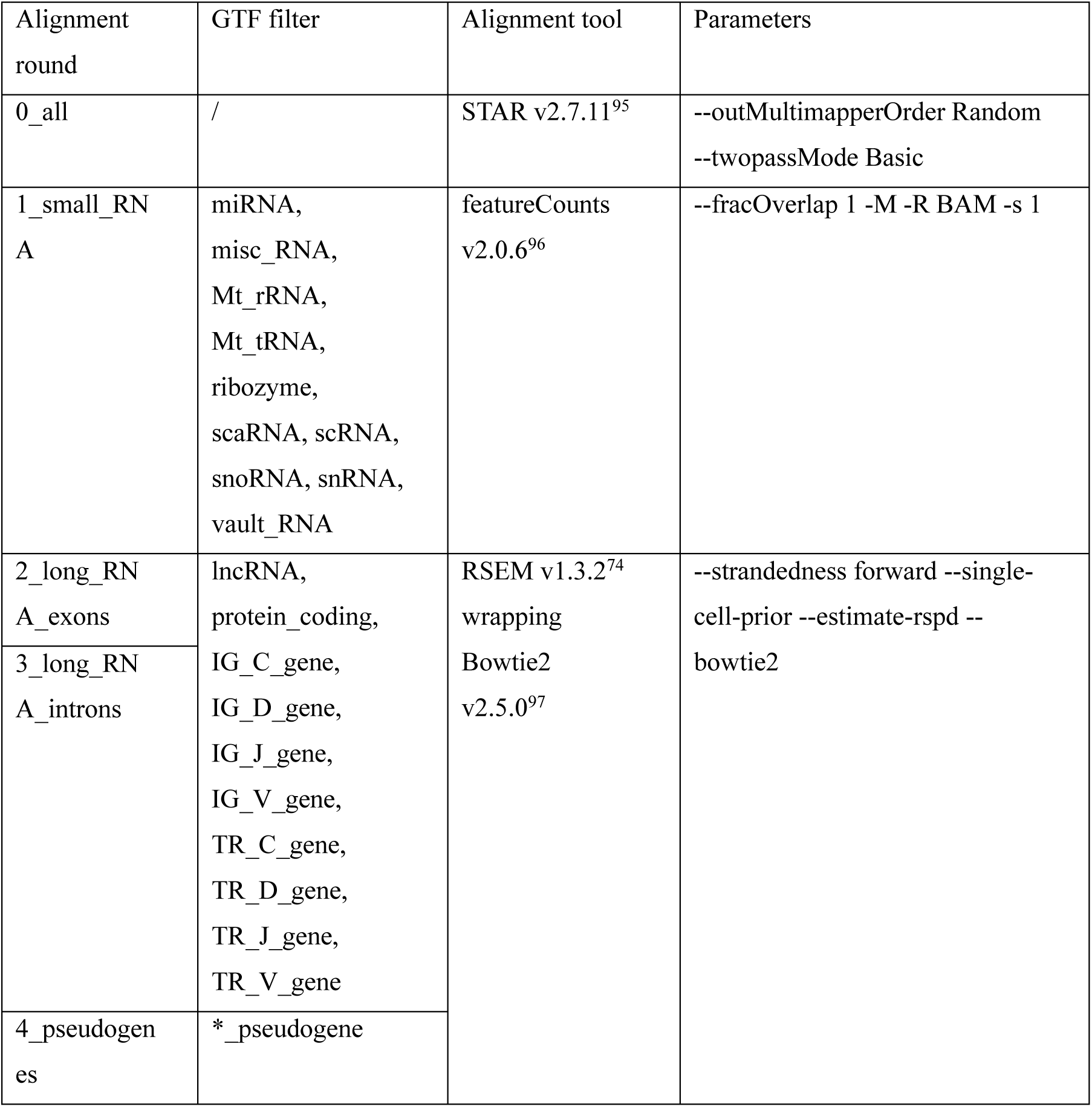
Alignment tools.

#### Single-cell total RNA expression analysis

The single-cell matrices from four alignment steps were loaded into R v4.4.1^98^ and combined for processing using the Seurat v 5.1.0^99^. Unnamed genes (gene symbols beginning with ENSG) were removed from the analysis for clarity. Data QC was processed using scCustomize v2.1.2^100^ to decide filtering thresholds, as listed in Table 3. For naïve PC9 inDrop and sc-rDSeq data, the normalization, scaling, and variable feature selection were performed using default settings. Principal component analysis (PCA) was conducted on the scaled data, with the number of principal components listed in Table 3. Clusters of cells were identified using a shared nearest neighbor (SNN) modularity optimization-based clustering algorithm, with resolutions specified in Table 3. UMAP was used for visualization.

**Table 3.** Seurat parameters for single cell gene expressions.

| Experiment | Threshold | PCs | resolution |
| --- | --- | --- | --- |
| PC9 sc-rDSeq | 1000 < nGene; nUMI < 250000 | 1:8 | 0.5 |
| PC9 inDrop | 1000 < nGene; nUMI < 20000 | 1:8 | 1 |
| PC9 + Drug<br>(gene) | 400 < nGene < 7000; 1500 < nUMI < 100000 | 1:10 | 1 |
| PC9 + Drug<br>(Isoform) | 300 < nGene < 5000; 1500 < nUMI < 80000 | 1:20 | 1 |

For PC9 Drug treatment data, the data analysis was split into two stages. Firstly, all data were analyzed with integration to remove technical noises using Harmony v1.2.3^101^. Then cell cycle phase scores of each cell were calculated based on canonical markers^102^ with Seurat function CellCycleScoring. Histone mRNA were split into replication dependent and independent^40^. Replication dependent histone mRNA scores were added to single cell and plotted in histogram with density plot. A normalized distribution curve is set up by setting 𝜇 as the peak of the density, and 𝜎 from the left distribution width. Cells with module score greater than 𝜇 + 2 × 𝜎^2^ are defined as S Phase while the rest are Non-S phase. Then only DTP were subclustered. Due to the big variance of coverage, first a harsh filter is applied to only keep high-quality cells, variable feature selection were selected using default settings, and kept as highly-variable genes for the next round of analysis, where a normal filter is applied to keep most of the cells (Table 3), and PCA analysis was performed with the highly-variable genes defined previously. Clusters of cells were identified using a shared nearest neighbor (SNN) modularity optimization-based clustering algorithm, with resolutions specified in Table 3. UMAP was used for visualization.

#### Gene-set enrichment analysis

Gene-set enrichment analysis was performed with ShinyGO v8.0^103^ and clusterProfiler v4.12.6^104^. Marker genes for each cluster were selected using Seurat’s FindAllMarkers function, applying a filter for a log2 fold change greater than 0 and an adjusted p-value less than 0.05. Customized background genes for the ShinyGO analysis included all expressed genes in the dataset. Marker genes for each cluster were mapped against the MSigDB Hallmark and GO Biological Processes pathway databases, using a filter with an FDR q-value less than 0.05.

#### Copy Number Variation analysis

The CNV analysis was conducted using InferCNV^62^ v3.2 using default parameters, with cells treated with DMSO served as the reference group, representing a normal karyotype, while cells treated with Osimertinib and Gefitinib constituted the observation groups, potentially harboring CNVs. Human genome (GRCh38, Ensembl Release 112) was used for gene location information.

#### eRNA analysis

Enhancer RNAs are found by searching for regions with enriched bidirectional peaks on the genome. sc-rDSeq data were realigned to the genome (GRCh38, Ensembl Release 112) using Bowtie 2, as STAR alignment results contain junction information that can confound the peak calling process. Alignments from each cluster, as defined by gene expression analysis in Seurat, were merged to identify enriched peak regions using HOMER v5^105^ findPeaks with parameters -style histone -region -size 150 -minDist 370 -F 0 -L 0 -C 0. The directionality of the peaks was calculated by dividing the number of reads from the plus strand by the total reads of the peak. Peaks with a directionality in the range of 5% to 95% were defined as bidirectional. Peaks covering more than two genes in opposite directions were removed. Additionally, peaks with low expression (<10 reads per peak) were also filtered out. Bidirectional peaks from all clusters were merged to form a list of candidate enhancer RNAs. These candidate eRNAs were characterized using HOMER annotation. The expression of eRNA in each cluster was determined by intersecting the candidate eRNA list with the merged alignment from each cluster, then normalized by read depth and scaled by row. The scaled expression matrix was clustered and reordered using AgglomerativeClustering from the Scikit-learn package^106^ with the number of clusters set to match the input. Candidate eRNAs were then intersected with eRNAs defined by the GeneHancer database^67^ and the connected gene list for each eRNA was retrieved. For each gene expression cluster, differentially upregulated genes (avg_log2FC > 0, p_val_adj < 0.05) were compared with the genes connected to eRNA expression clusters. Overlapping genes were identified as those upregulated in both gene expression and eRNA regulation. Whether the connected genes found in the list of differentially expressed genes were due to chance was tested. First, same number of eRNAs as the input eRNA per cluster were randomly sampled from GeneHancer. Then it was tested whether the number of overlap genes between cluster’s DE genes and the candidate eRNA connected genes was significantly greater than the number of overlap genes between cluster’s differentially expressed genes and random eRNA connected genes. The test was conducted by performing 50 sampling iterations to obtain a p-value, and this process was repeated ten times for the distribution of p-values.

#### Isoform analysis

The AS events and junction counts were analyzed by rMATS^107^ using default parameters for naïve PC9 data of inDrops and sc-rDSeq. The isoform analysis was performed on pseudo-bulk of cellular clusters. DTP cells were grouped according to the single cell clustering annotations, raw fastq files were merged as pseudo-bulk. For each cluster, the isoform calculation is based on the comparison between individual clusters against all the rest of the cells as background. In each dataset, three samples were randomly picked from all reads, with sampling size equal 70% of the number of reads of the pseudo bulk cluster. Then alignment for transcriptome (not including introns) using RSEM with Estimation Maximization (EM) algorithm was performed to get isoform counts. The expected counts were then used for isoform switch analysis with IsoformSwitchAnalyzeR^108^ v2.4.0. In brief, isoform counts were imported and normalized^109^, then isoform switch test was performed^110,111^. Significantly switched isoforms were also checked for protein domains using Pfam^112^ and signaling peptides with SignaIP^113^ v6.0.

Single cell isoform analysis was performed on isoform matrices obtained on single cell RSEM alignment. Data QC was processed using scCustomize v2.1.2^100^ to decide filtering thresholds, as listed in Table 3. After filtering and normalization as default, top 2000 switched isoforms retried from the pseudo-bulk isoform switch analysis were used for data scaling and PCA. Clusters of cells were identified using a shared nearest neighbor (SNN) modularity optimization-based clustering algorithm, with resolutions specified in Table 3. UMAP was used for visualization. Then gene expression based cell cluster annotations were compared with that of isoform expression, fraction of cells that overlaps between each pair of clusters were calculated and presented in a heatmap.

#### SNVs and indels analysis

Detection of mutations was performed on treatment-level pseudo-bulks using GATK^79^ v4.5.0.0 Mutect2. Alignments of single cells from each condition were merged into condition pseudo bulks, then transferred to Mutect2 in tumor-only mode to call mutations. Gefitinib and Osimertinib mutation calling was done with default parameters, DMSO specific parameters were: --initial-tumor-lod 1 --tumor-lod-to-emit 1 --max-population-af 1 to allow detection of very rare instances that could potentially give rise to DTP cells. The following step was GATK v4.5.0.0 FilterMutectCalls, and eventually we filtered out sites with filters defined as FAIL, weak_evidence, map_qual, base_qual, and contamination. Additionally, sites were annotated with GATK v4.5.0.0 Funcotator with gencode source. We filtered any sites overlapping with known RNA-editing sites^114–116^, only sites that were covered with at least one read in all 3 conditions were kept, and eventually only sites with at least 10 reads at either Gefitinib or Osimertinib were taken for testing enrichment/depletion compared to DMSO. Each site was tested with chi-square or fisher-exact depending on sample sizes for the allele frequencies, then FDR correction was applied and threshold for significance was defined as 0.05. Sites with variant classification of missense and splice-site were kept for further observation. We applied SComatic^87^ SingleCellGenotype on DTP cells in these candidate sites with --min_mq 20.

To test the EGFR mutational profiles across the conditions we extracted EGFR-annotated sites after filtration of FAIL, weak_evidence, map_qual, base_qual, and contamination. We used DP (depth) and AF (allele frequency) to calculate the number of mutated reads. To measure overlap between mutations in each condition we used both genomic location and substitution information.

## Supplemental information

File S1. Supplementary File 1.sc-rDSeq cost calculation.xlsx. Related to Fig.1

File S2. Supplementary File 2.sc-rDSeq Barcode primers.zip. Related to methods.

---

**Fig S1.**
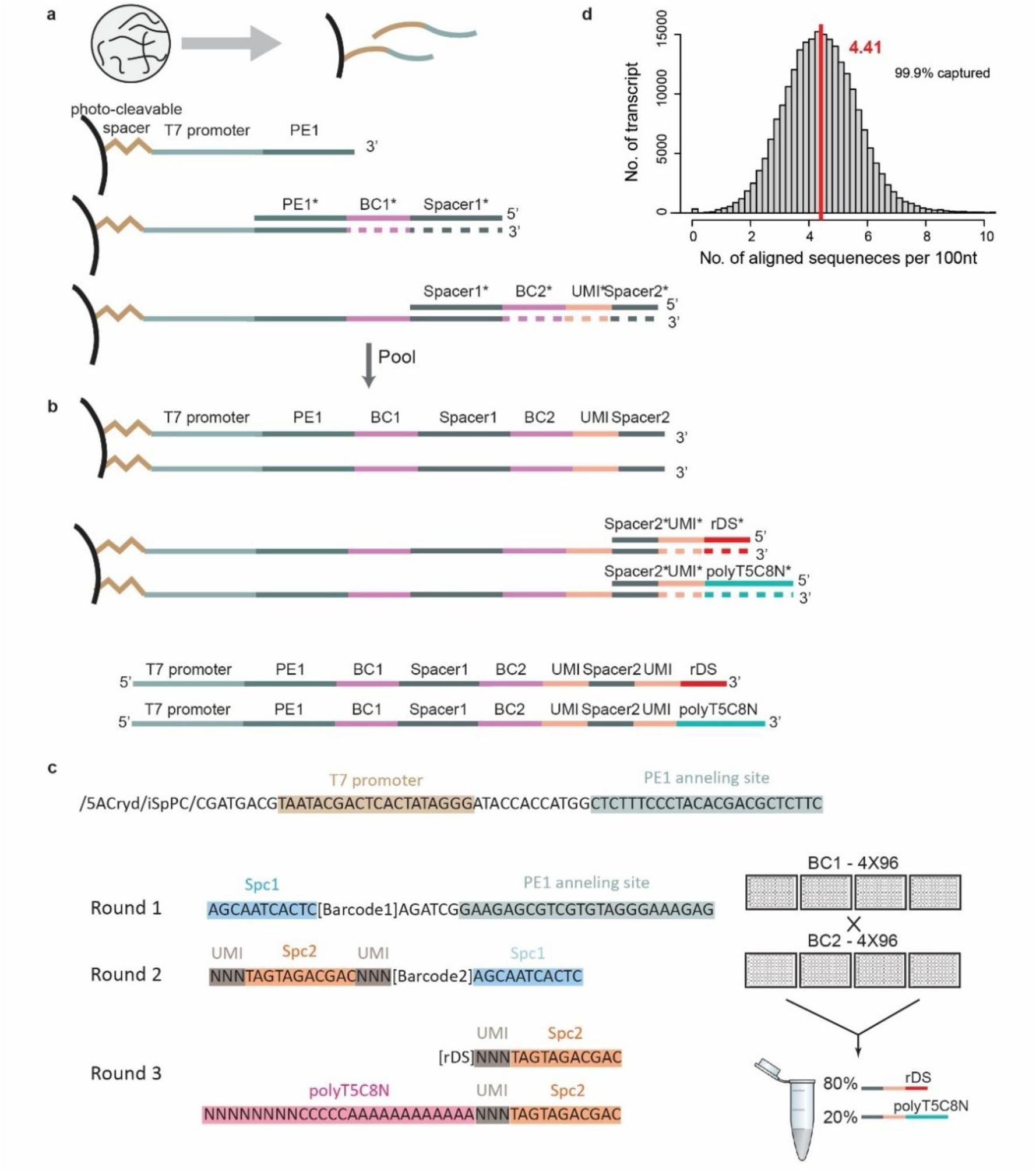
sc-rDSeq barcode design and fabrication. **a)** Barcoding hydrogel beads are synthesized by incorporating acrydite primers. Barcode diversity increases geometrically through a ‘split-and-pool’ strategy. **b)** Beads are pooled for rDS and polyT5C8N primer extension at an 8:2 ratio, ensuring each bead shares BC1 and BC2, with 80% rDS and 20% polyT5C8N. **c)** Primer sequences used in each round. The schema on the right illustrates single-cell barcode complexity, resulting from two primer sets distributed across four 96-well plates and later combined for rDS and polyT5C8N extension. **d)** *in silico* rDS sequence alignment to the transcriptome, with a histogram showing primer annealing frequency per 100 nt. The red line and number indicate the median primers aligned, while the top-left percentage represents the proportion of captured transcripts.

**Fig S2.**
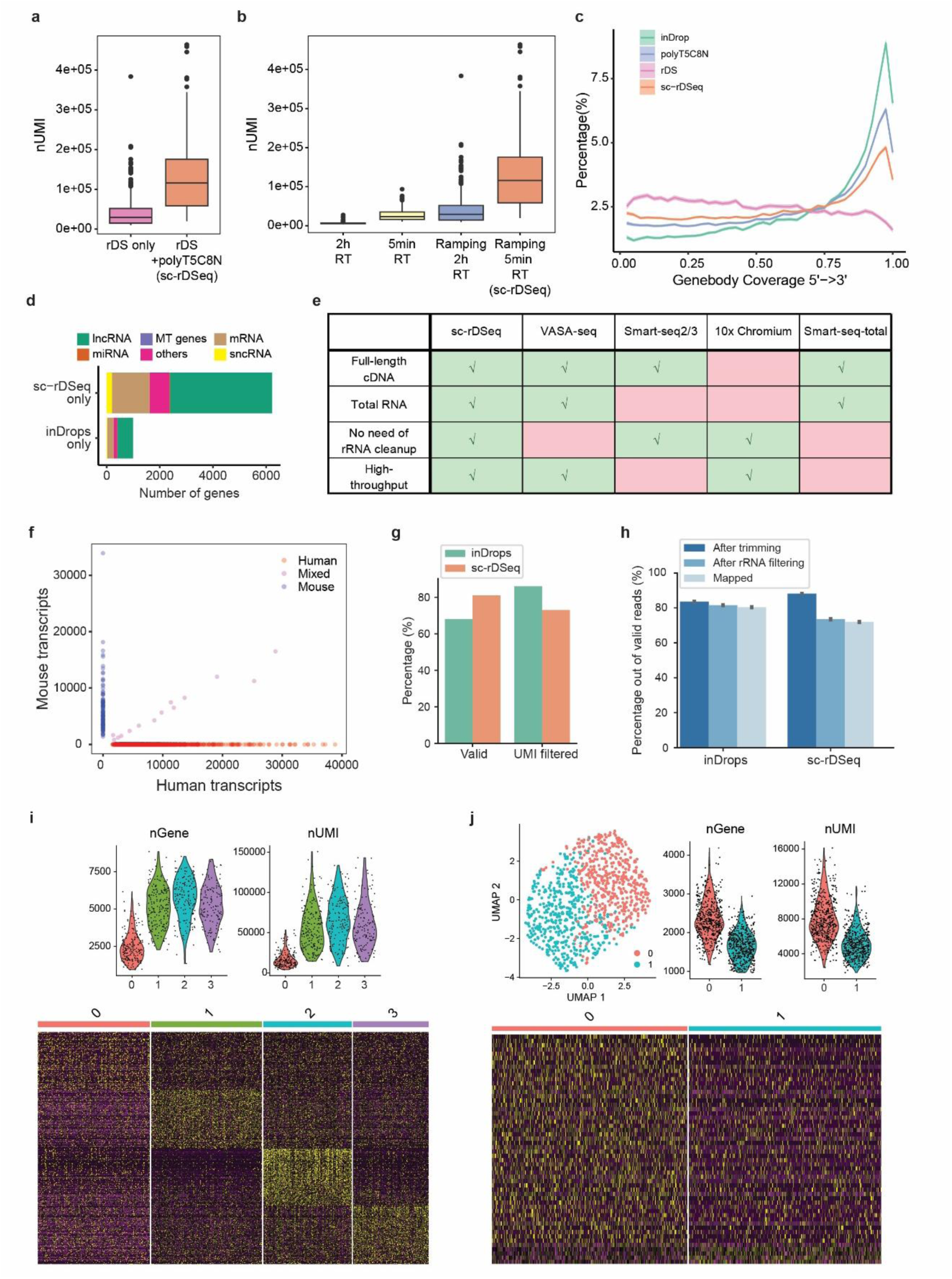
sc-rDSeq quality control. **a)** nUMI per cell with different RT strategies, of continues RT reaction (50°C) for 2h or 5min, or gradually ramp from 4°C to 50°and repeat for 20 cycles allowing total time at 50°C for 2h or 5min. Mean from left to right: 5993, 23313, 29464.5, 116117. **b)** nUMI per cell with different primer sets, with only rDS primers and combination of rDS and polyT5C8N primer in 8:2 ratio. Mean from left to right: 29464.5, 116117. **c)** Gene-body coverage across all genes based on merged single cells data from inDrops, sc-rDSeq, and sc-rDSeq split into reads originated from rDS primers and polyTN8 primers. Each gene is divided into 40 segments and counts failing into each segment are calculated for coverage. Ribbon area shows the SEM around mean. **d)** Number of genes by each type of the RNAs uniquely detected by sc-rDSeq or inDrops. **e)** Technical parameters comparing sc-rDSeq with other single cell full-length RNA seq technologies. **f)** sc-rDSeq of human and mouse cells. The scatter plot shows the number of transcripts associated to each barcode. Blue and red dots indicate human- and mouse-specific transcripts, while the purple dot indicates a mixed association. **g)** Out of the total raw reads, the percentage of reads have correct library structure (valid) and out of which have unique UMI (UMI filtered), comparing inDrops and sc-rDSeq. **h)** Single cell reads percentage remained after each round of data pre-processing, including adaptor trimming, rRNA filtering and genome mapping. **i)** sc-rDSeq data of PC9 cells with violin plots showing coverage by number of genes and UMIs per cell in each cluster defined by sc-rDSeq analysis for PC9 cell (top), differentially expressed genes were identified using KNN unsupervised clustering of total transcriptome expression and presented in heatmap of gene expression (bottom). **j)** inDrops data of PC9 cells with UMAP analysis (n = 1193; top left) and violin plots showing coverage by number of genes and UMIs per cell (top right), and no heterogeneity observed between the two clusters obtained based on transcriptome expression as shown in heatmap of gene expression (bottom).

**Fig S3.**
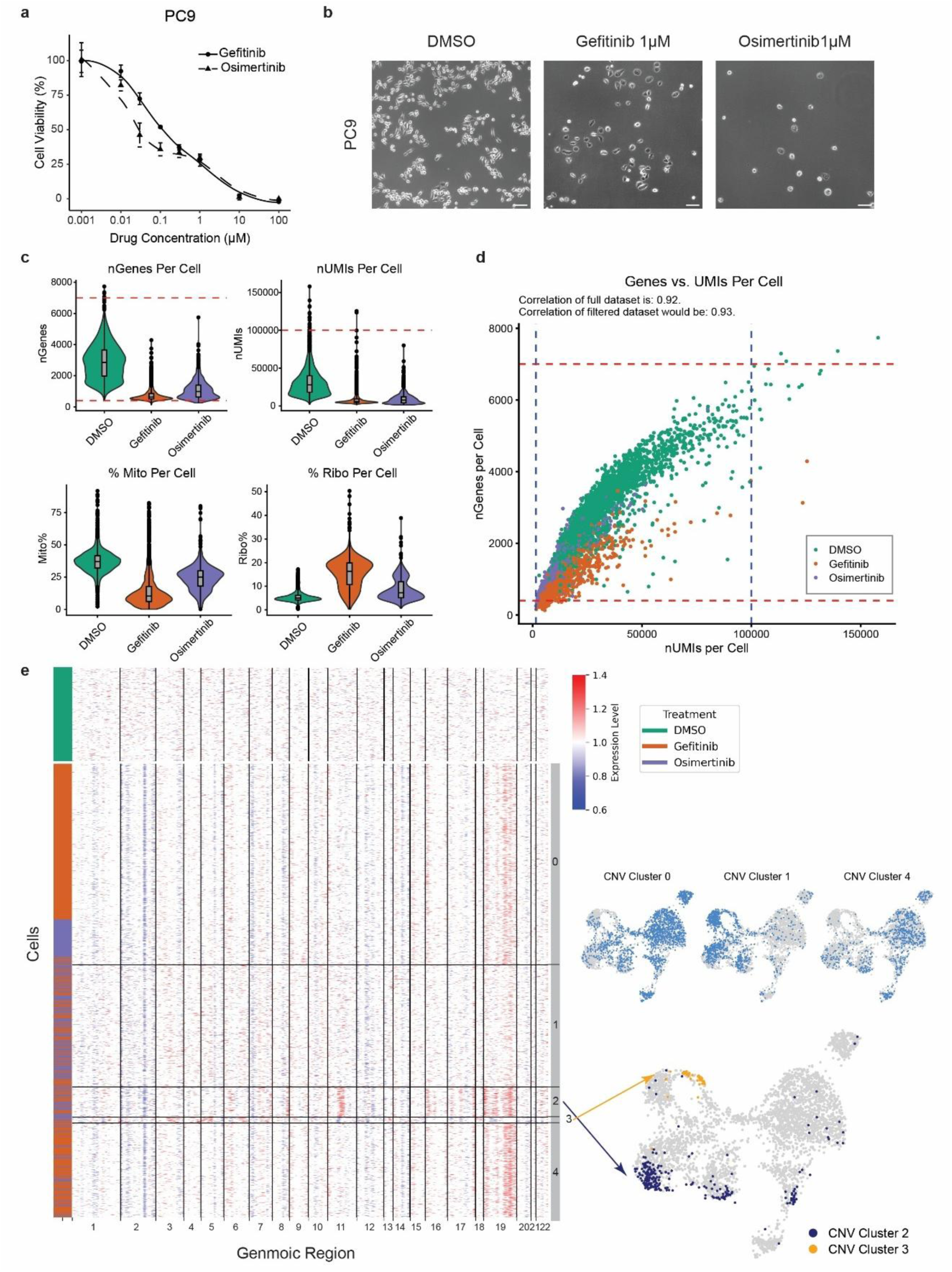
Drug persistent experiments and quality control. **a)** Dose response curve of NSCLC cell lines: PC9 in response to two EGFR inhibitors: Gefitinib and Osimertinib. Gefitinib IC50 = 118 nM, Osimertinib IC50 = 122 nM. **b)** Microscopy photos of PC9 cells under treatment of DMSO for 3 days, 1 μM Gefitinib and 1 μM of Osimertinib for 9 days. Scale bar: 100 μm. **c)** QC showing number of genes, UMIs and mitochondrial and ribosomal RNA percentage per cell, divided by groups. Red dot lines highlight the threshold for data filtering. **d)** Correlation between nGenes and nUMIs per cell, red and blue dot lines highlight the threshold for data filtering. **e)** InferCNV analysis of DTPs with untreated cells as references. Heatmap clustered using agglomerative clustering. Cells from each CNV cluster were highlighted separately on gene expression UMAP.

**Fig S4.**
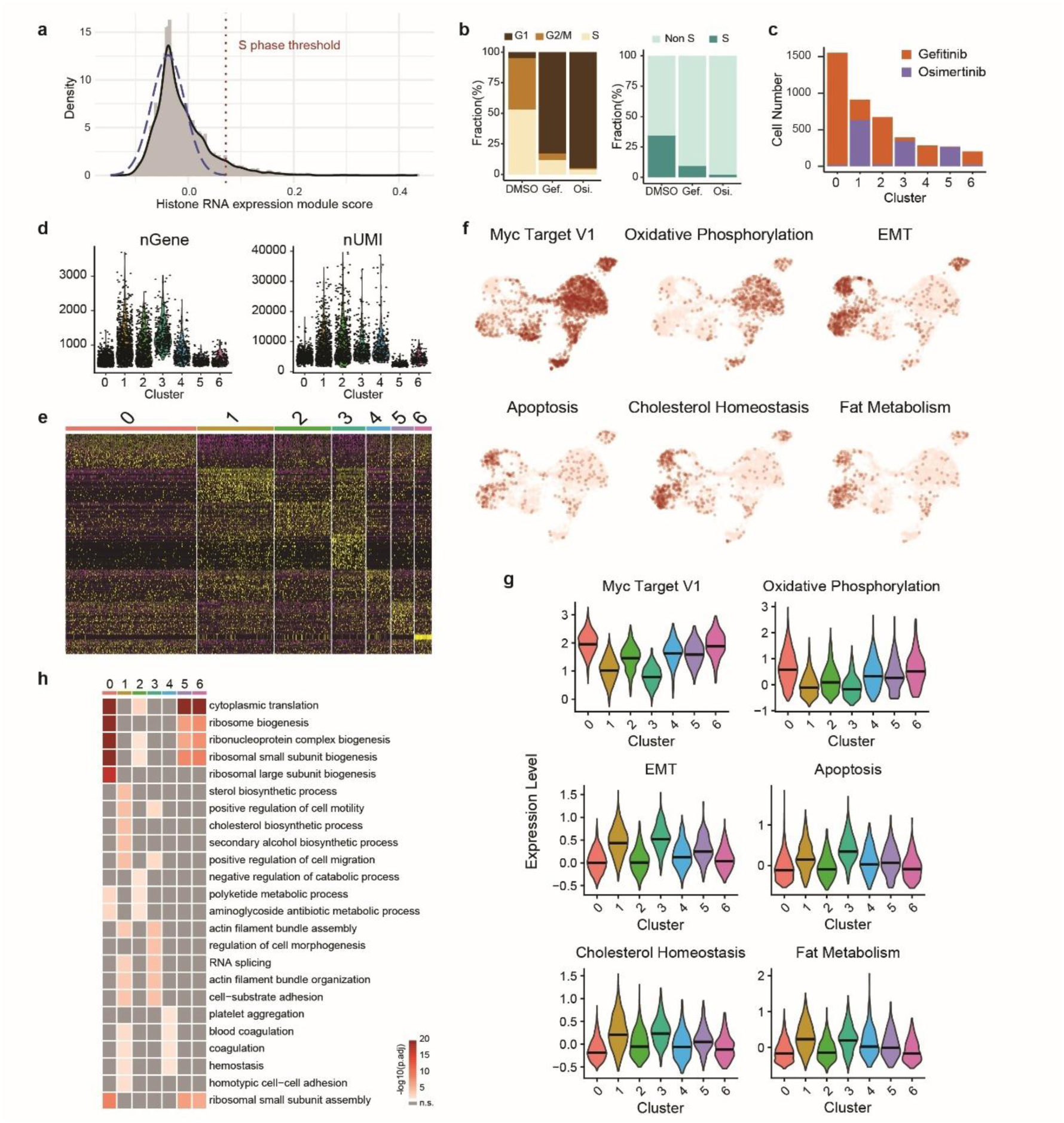
sc-rDSeq reveals heterogeneity based on total RNA in PC9 drug tolerant persisters. **a)** Histogram with density plot showing replication-dependent histone mRNA expression scores per single cell, displaying a normalized distribution with a right-skewed tail. The threshold for defining S and non-S phase cells is set at the mean plus two standard deviations. **b)** Percentage of cells in each phase of cell cycle by treatment, defined by cell cycle marker gene expression for G1, G2/M and S phases (left), or histone mRNA expression for S and non-S phases (right). **c)** Number of cells by treatment in each cluster. Mapping of clusters to their major contributor of treatment: Clusters 0, 2, 4, and 6 – Gefitinib; Clusters 1, 3, and 5 – Osimertinib. **d)** Number of genes and UMIs per cluster of cells. **e)** Heatmap of differentially expressed genes identified via KNN unsupervised clustering. **f)** Feature plots showing module scores of upregulated pathways. **g)** Violin plot showing pathway module scores by clusters. **h)** Heatmap of significantly upregulated GO BP pathways per clusters including genes for ribosomal proteins.

**Fig S5.**
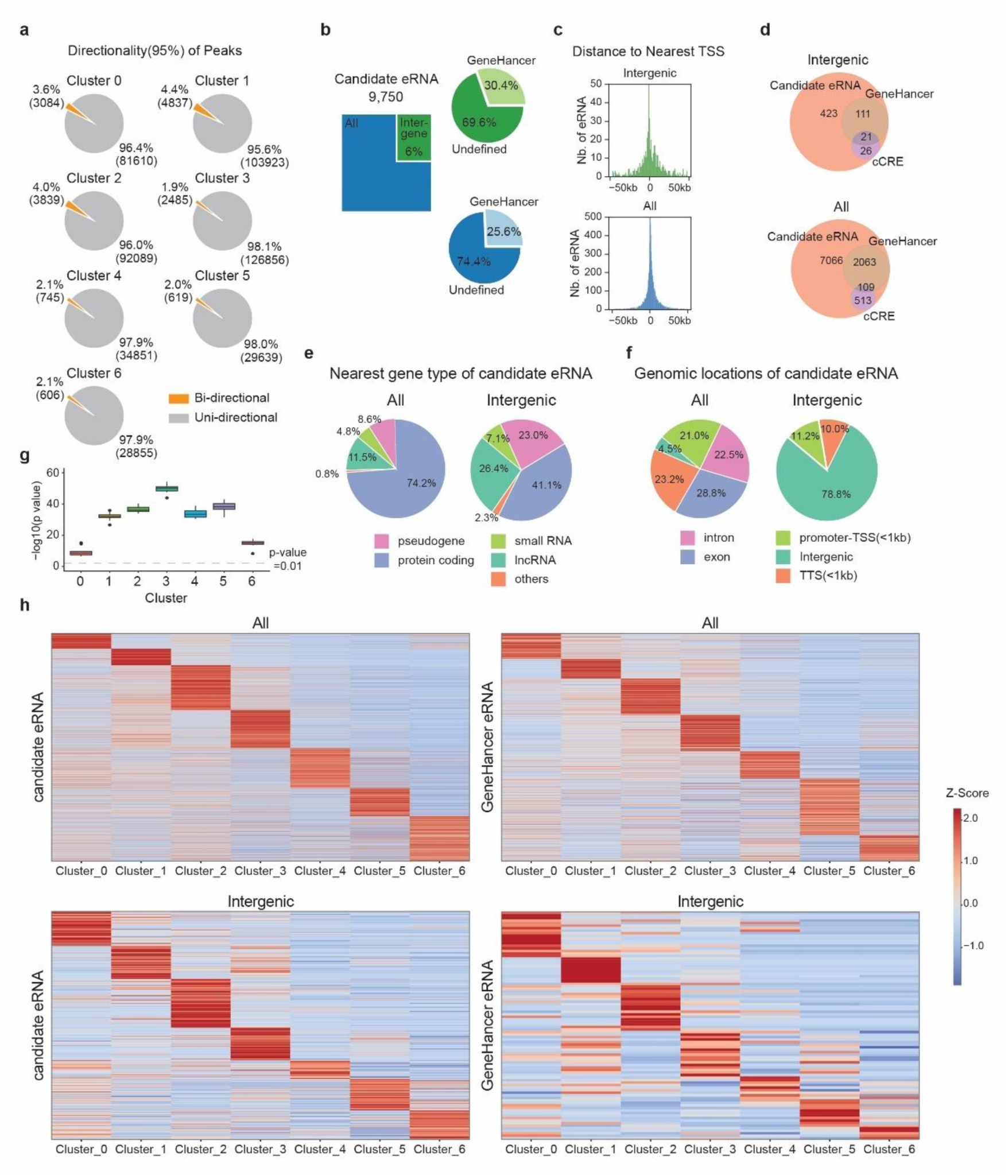
eRNA expression is correlated with gene expression. **a)** Directionality of peaks per cluster. Peaks are defined as bidirectional if less than 95% of the reads come from the same strand. **b)** Number of candidate eRNAs (left) with fractions that overlap with GeneHancer (right), distinguishing between intergenic (top) and all (bottom) eRNAs. **c)** Distribution of candidate eRNAs by the nearest transcription start site (TSS), distinguishing between intergenic (top) and all (bottom) eRNAs. **d)** Overlaps of candidate eRNAs with two eRNA databases, GeneHancer and cCRE, distinguishing between intergenic (top) and all (bottom) eRNAs. **e)** and **f)** Distribution of the types of nearest genes (e) and the genomic locations (f) to candidate eRNAs, distinguishing between intergenic (top) and all (bottom) eRNAs. **g)** Boxplot displaying p-values derived from a shuffling t-test, which evaluates the enrichment of candidate eRNA-related genes in differentially expressed genes compared to random enhancers sampled from GeneHancer, for each cluster. **h)** Heatmaps depicting expression levels of candidate eRNAs in pseudo-bulk clusters defined by gene expression UMAP clustering. The heatmaps are stratified by genomic location (all eRNAs vs. intergenic eRNAs, top and bottom panels) and their presence in the GeneHancer database (right and left panels). Expression levels were normalized by read depth per cluster and centered by row around zero, represented as z-scores. Rows are ordered according to agglomerative clustering results.

**Fig S6.**
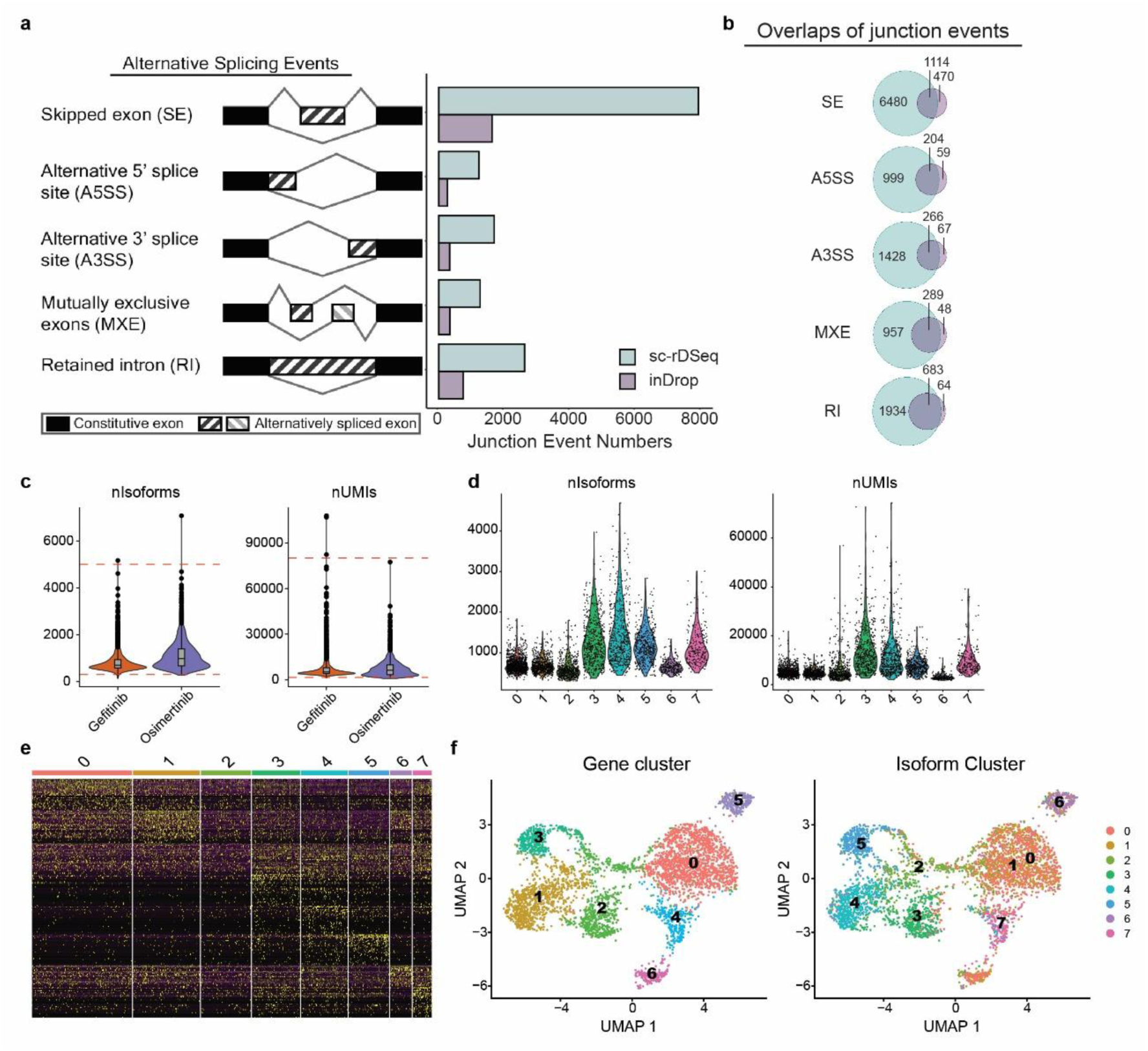
Isoform detection by sc-rDSeq. **a)** and **b)** Number of alternative splicing events (a) and junction events (b) detected in PC9 cells using sc-rDSeq and inDrops. Alternative splicing events quantified by rMATS. **c)** Number of Isoforms and UMIs per single cell by group. Red dot lines highlight the threshold for data filtering. **d)** Number of isoforms and UMIs per single cell by isoform expression clusters. **e)** Heatmap of differentially expressed isoforms per cluster. **f)** UMAP representation of gene expression profile annotated by gene expression defined clusters (left) and isoform expression defined clusters (right).

**Fig S7.**
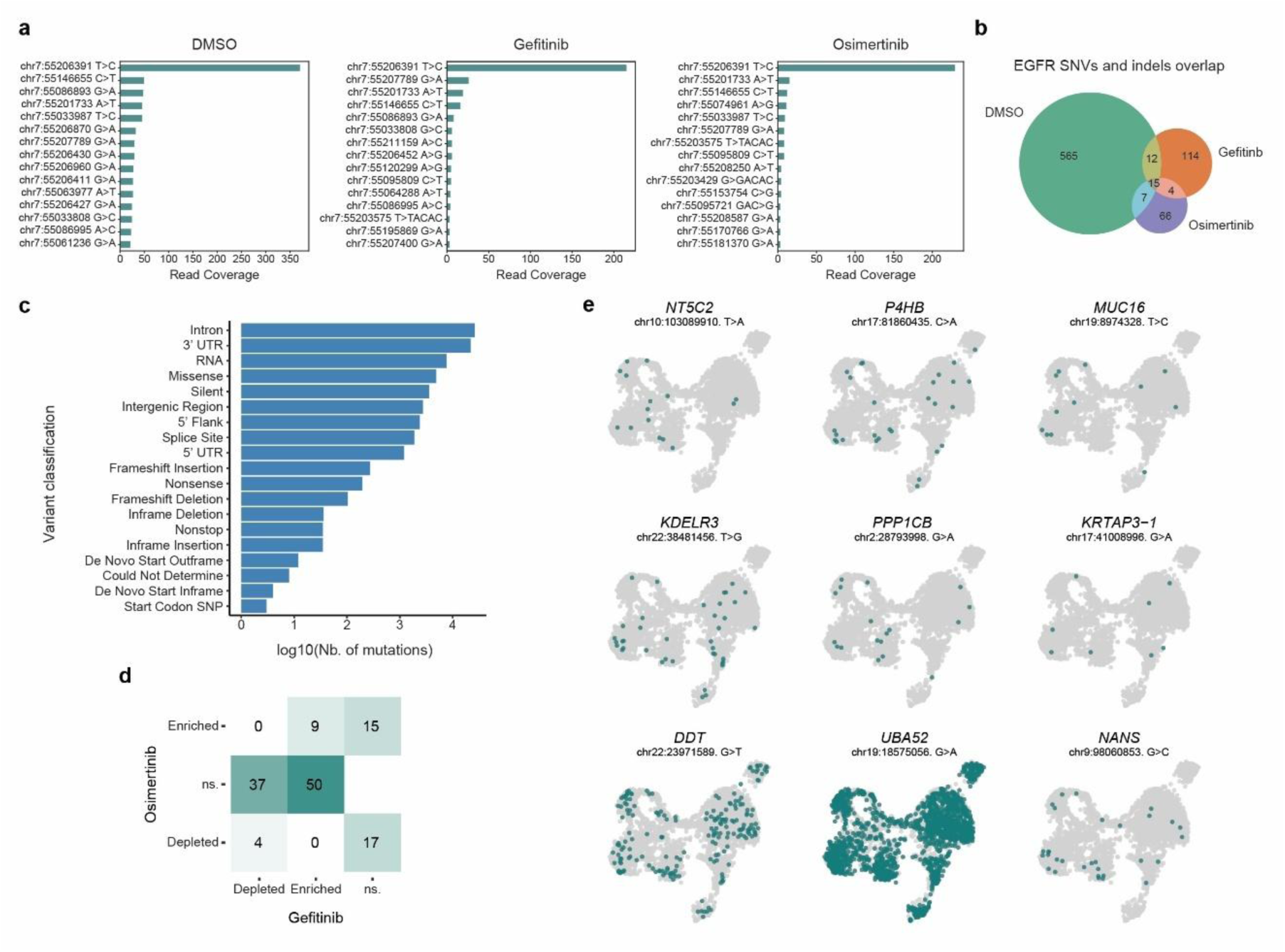
Genetic variation analysis in reveals non-driver role in DTP heterogeneity. EGFR SNVs and indels in DMSO, Gefitinib and Osimertinib DTPs with **a)** mutation read coverage of the top 15 SNVs and indels and **b)** overlaps between groups. **c)** Variant classification counts of all identified mutation. **d)** Number of enriched and depleted SNVs intersect between Gefitinib- and Osimertinib-DTPs. **e)** UMAP visualization highlighting cells carrying selected mutations.

